# Exon junction complex-associated multi-adapter RNPS1 nucleates splicing regulatory complexes to maintain transcriptome surveillance

**DOI:** 10.1101/2021.08.20.457088

**Authors:** Lena P. Schlautmann, Volker Boehm, Jan-Wilm Lackmann, Janine Altmüller, Christoph Dieterich, Niels H. Gehring

**Affiliations:** Institute for Genetics, University of Cologne, 50674 Cologne, Germany; Center for Molecular Medicine Cologne (CMMC), University of Cologne, 50937 Cologne, Germany; CECAD Research Center, University of Cologne, Joseph-Stelzmann-Str. 26, 50931 Cologne, Germany; Cologne Center for Genomics (CCG), University of Cologne, 50931 Cologne, Germany; Berlin Institute of Health at Charité – Universitätsmedizin Berlin, Core Facility Genomics, Charitéplatz 1, 10117 Berlin, Germany and Max Delbrück Center for Molecular Medicine in the Helmholtz Association (MDC), Berlin, Germany; Section of Bioinformatics and Systems Cardiology, Department of Internal Medicine III and Klaus Tschira Institute for Integrative Computational Cardiology, Heidelberg University Hospital, 69120 Heidelberg, Germany; DZHK (German Centre for Cardiovascular Research), Partner site Heidelberg/Mannheim, 69120 Heidelberg, Germany

**Author notes:** Contact: Niels H. Gehring.

## Abstract

The exon junction complex (EJC) is an RNA-binding multi-protein complex with critical functions in post-transcriptional gene regulation. It is deposited on the mRNA during splicing and regulates diverse processes including pre-mRNA splicing, mRNA export, mRNA translation, and nonsense-mediated mRNA decay (NMD) via various interacting peripheral proteins. The EJC-binding protein RNPS1 might serve two functions: it suppresses mis-splicing of cryptic splice sites and activates NMD in the cytoplasm. When analyzing the transcriptome-wide effects of EJC and RNPS1 knockdowns in different human cell lines, we find no evidence for RNPS1 being a globally essential NMD factor. However, various aberrant splicing events strongly suggest that the main function of RNPS1 is splicing regulation. Rescue analyses revealed that about half of these RNPS1-dependent splicing events was fully or partially rescued by the expression of the isolated RRM domain of RNPS1, whereas other splicing events are regulated by its C-terminal domain. We identified many splicing-regulatory factors, including SR proteins and U1 snRNP components, that specifically interact with the C-terminus or with the RRM of RNPS1. Thus, RNPS1 emerges as a multifunctional splicing regulator that promotes correct and efficient splicing of different vulnerable splicing events via the formation of diverse splicing-promoting complexes.

## Introduction

The majority of human genes contain introns and their transcribed pre-mRNAs are subject to (alternative) splicing (1). During splicing, intronic sequences are excised and exons are ligated by the spliceosome, resulting in a mature mRNA that is subsequently exported to the cytoplasm (2). The spliceosome has the critical as well as delicate task to identify the correct splice sites, because frequently there is more than one possible splice site. These additional sites can be designated alternative splice sites, but also so-called cryptic splice sites (3). While the usage of the former can be employed to generate different transcript isoforms, the erroneous utilization of cryptic splice sites leads to mis-splicing and the production of defective transcripts (4). Both processes use similar mechanisms, but with opposing results. Alternative splicing (AS) increases the number of protein isoforms produced from a single gene by the varying usage of 5’ and 3’ splice sites, skipping of exons and inclusion of introns (5). In addition, it is also an important step in gene expression regulation. In contrast, the use of cryptic splice sites is often associated with the production of non-functional transcripts and the occurrence of disease (4). Therefore, (alternative) splicing requires tight regulation and accurate operation of the spliceosome to ensure the production of the correct mature mRNAs.

The recognition of the proper splice sites by the spliceosome is assisted by auxiliary proteins, which bind to the pre-mRNA and guide the spliceosome to the correct positions (6). The group of splicing regulatory proteins is quite diverse and includes several RNA-binding proteins that interact with specific sequence motifs, such as the SR proteins (7). The class of SR proteins is characterized by one or two N-terminal RNA binding domains (e.g. RRMs), and a C-terminal domain enriched in arginine and serine dipeptides (RS-domain). SR proteins bind to exonic splicing enhancers (ESEs) and thereby define the exons to be maintained in the mature mRNA (8). In contrast, hnRNPs bind mainly to intronic sequences (intronic splicing enhancers, ISEs) and support their recognition and removal by the spliceosome (9,10). The loss or exchange of ESE and ISE sequences can have dramatic effects on the splicing pattern and might even result in the inactivation of genes (11). However, the splicing process is not only regulated by SR proteins and hnRNPs, but also by many other RNA-binding proteins (12). Among these is also the exon junction complex (EJC), an RNA-binding protein complex, which binds 24 nt upstream of an emerging exon-exon junction, independent of the RNA sequence (13). The EJC core consists of the three proteins EIF4A3, RBM8A and MAGOH that are deposited onto the mRNA during splicing by interactions of the EJC proteins with spliceosome components (14,15).

The EJC carries out diverse functions during post-transcriptional gene regulation and besides regulating pre-mRNA splicing, it also facilitates the transport and translation of spliced mRNAs (13). Furthermore, the EJC is critical for nonsense-mediated mRNA decay (NMD) and EJC proteins were initially described to enable the detection of NMD substrates, particularly mRNAs containing premature translation termination codons (PTCs) (16,17). NMD not only serves as a quality control mechanism by ensuring the degradation of incorrect mRNAs, but is also important for the regulation of gene expression (18). NMD substrates can be produced in different ways: NMD-activating termination codons may result from AS or genomic mutations, in other cases NMD is triggered by a long 3’ UTR. Efficient NMD takes place when a ribosome terminates at a PTC. EJCs bound to the mRNA downstream of that PTC will serve as a signal for the NMD machinery to initiate the degradation of that mRNA. Therefore, the EJC is essential for NMD to correctly identify transcripts that need to be degraded.

In order to carry out all its different tasks, the EJC functions as a binding platform for auxiliary factors which itself have varying regulatory potentials. Two of the EJC-associated complexes, the apoptosis and splicing associated protein complex (ASAP) and the PSAP complex are known regulators of splicing (19). They share two of their components, RNA binding protein with serine rich domain 1 (RNPS1) and Sin3A associated protein 18 (SAP18), but vary in the third component, which is either Acinus (ACIN1) or Pinin (PNN), respectively (20,21). A previous study has shown that knockdown (KD) of PNN and ACIN1 affects different splicing events, suggesting that ASAP and PSAP complexes have non-overlapping functions in splicing regulation (22). In *D. melanogaster*, retention of PIWI intron 4 relies on the EJC core, ACIN1 and RNPS1 (23,24). Recent studies furthermore demonstrate the ability of the EJC core and the PSAP to suppress the usage of cryptic 5’ and 3’ splice sites (25-28). While cryptic 3’ splice sites are suppressed by direct masking by the EJC core, the suppression of cryptic 5’ splice sites involves an unknown mechanism requiring RNPS1 recruitment via the EJC core and the PSAP complex (25,26). It is likely that RNPS1 represents the central functional component in all these processes, whereas ACIN1 and probably PNN as well play a role in RNPS1 recruitment. ACIN1, for example, directly binds to the EJC core (22), which would explain how the interaction of RNPS1 and the EJC is established.

In addition to its function in splicing, RNPS1 also has the ability to activate NMD when tethered to a reporter mRNA downstream of the termination codon (17,29). It has also been reported that the presence of RNPS1 on NMD-targeted mRNAs leads to more pronounced degradation (30). However, there are controversial results as to whether RNPS1 has an essential role in NMD or not (31,32). Although the exact function of RNPS1 during NMD remains to be determined, it is clear that RNPS1 interacts with the EJC and possibly also with components of the NMD machinery, potentially forming a bridge between these two macromolecular assemblies.

Although previous work had examined individual aspects of RNPS1, its function in the context of the EJC is still not fully understood and therefore demands a more comprehensive characterization of RNPS1. In this study, we uncover that RNPS1 only mildly affects a small subset of NMD targets and its main function is the regulation of AS. To that end, the RNPS1 RRM, which is known to be required for ASAP/PSAP assembly, regulates splicing by binding other splicing factors, including SR proteins and spliceosomal components (20). We identified many components of the U1 snRNP that interact with the C-terminus and thus conclude that RNPS1 is a part of and bridges different splicing competent complexes to the EJC to regulate splicing of surrounding/adjacent introns. In our model RNPS1 acts as a multi-functional adapter that recruits splicing factors independently of the mRNA sequence to the EJC binding site.

## Material and Methods

### Cell Culture

Flp-In-T-REx-293 (HEK 293; human, female, embryonic kidney, epithelial; Thermo Fisher Scientific, RRID:CVCL_U427), HeLa Flp-In-T-REx (HeLa FT; human, female, cervix; Elena Dobrikova and Matthias Gromeier, Duke University Medical Center) and HeLa Tet-Off (HTO; human, female, cervix; Clontech, RRID: CVCL_V352) cells were cultured in high-glucose, GlutaMAX DMEM (Gibco) supplemented with 9% fetal bovine serum (Gibco) and 1x Penicillin Streptomycin (Gibco). The cells were cultivated at 37°C and 5% CO_2_ in a humidified incubator. The generation of stable cell lines is described below and all cell lines are summarized in Supplementary Table 1.

### Stable cell lines and plasmids

RNPS1 point and deletion mutants were PCR amplified using Q5 polymerase (New England Biolabs) and inserted into PB-CuO-MCS-BGH-EF1-CymR-Puro (modified from System Biosciences), together with an N-terminal FLAG-emGFP-tag via NheI and NotI (both New England Biolabs) restriction sites. As a control, FLAG-emGFP was equally cloned into the PB-CuO-MCS-BGH-EF1-CymR-Puro vector.

HEK 293 and HTO cells were stably transfected using the PiggyBac Transposon system. 2.5-3×10^5^ cells were seeded 24 h before transfection in 6-wells. 1 µg of PiggyBac construct was transfected together with 0.8 µg of the Super PiggyBac Transposase expressing vector using the calcium phosphate method. 48 h after transfection, the cells were transferred into 10 cm dishes and selected with 2 µg ml^-1^ puromycin (InvivoGen). After 7-10 days, the colonies were pooled. Expression of the PiggyBac constructs was induced with 30 µg ml^-1^ cumate.

RFX5 reporters were PCR amplified as described above and cloned into pcDNA5/FRT/TO/FLAG. HeLa FT cells were stably transfected with the reporters using the Flp-In-T-REx system. Transfection and selection was performed like for PiggyBac transfected cells, with the following differences: 1.5 µg of pcDNA5 construct were co-transfected with 1.5 µg Flippase expression vector (pOG44) and cells were selected with 100 µg ml^-1^ Hygromycin (InvivoGen). All cell lines generated and plasmids used in this study are listed in Supplementary Table 1.

### Co-Immunoprecipitation

Expression of FLAG-emGFP tagged RNPS1 mutants and FLAG-emGFP control was induced in stable cell lines (1.5 × 10^6^ cells per 10 cm dish) using cumate (as described above) 72 h before cell lysis. The samples were lysed in 600 µl buffer E (20 mM HEPES-KOH (pH 7.9), 100 mM KCl, 10% glycerol, 1 mM DTT, Protease Inhibitor) in the presence of 1 µg ml^-1^ RNase A and sonicated using the Bandelin Sonopuls mini20 with 10 pulses (2.5 mm tip, 1s pulse, 50% amplitude). For immunoprecipitation, the protein concentration of the lysates was measured using Pierce Detergent Compatible Bradford Assay Reagent (Thermo Fisher Scientific) and adjusted in buffer E. Then, the lysates were loaded onto Anti-FLAG M2 Magnetic Beads (Sigma-Aldrich) and incubated for 2 h at 4°C with overhead shaking. After that, the beads were washed four times for 3 min with mild wash buffer (20 mM HEPES-KOH (pH 7.9), 137 mM NaCl, 2 mM MgCl_2_, 0.2% Triton X-100, 0.1% NP-40). For elution, 2x 21.5 µl (42.5 µl total) of a 200 mg ml^-1^ dilution of FLAG peptides (Sigma) in 1x TBS was used.

### Label-free Mass Spec and computational analysis

For Label-free Mass spec, samples were immunoprecipitated as described above and after addition of 1 volume of 5% SDS in PBS reduced with DTT and alkylated with CAA (final concentrations 5 mM and 55 mM, respectively). For tryptic protein digestion, a modified version of the single pot solid phase-enhanced sample preparation (SP3) protocol was used as described below (33). Samples were reduced with 5 mM Dithiothreitol followed by alkylation using 40 mM Chloroacetamide. Afterwards, proteins were supplemented with paramagnetic Sera-Mag speed beads (Cytiva) and mixed in a 1:1-ratio with 100% acetonitrile (ACN). After 8 min incubation, protein-beads-complexes were captured using an in-house build magnetic rack, washed twice with 70% EtOH, and washed once with 100% ACN. After airdrying and reconstitution in 5 µl 50 mM triethylammonium bicarbonate, samples were supplemented with 0.5 µg trypsin and 0.5 µg LysC and incubated overnight at 37°C. The beads were resuspended on the next day and mixed with 200 µl ACN, followed by 8 min incubation. Subsequently, the samples were placed on the magnetic rack to wash the tryptic peptides once with 100% ACN. Samples were airdried, dissolved in 4% DMSO, transferred into new PCR tubes, and acidified with 1 µl of 10% formic acid. Proteomics analysis was performed by the proteomics core facility at CECAD via data-dependent acquisition using an Easy nLC1200 ultra high-performance liquid chromatography (UHPLC) system connected via nano electrospray ionization to a Q Exactive Plus instrument (all Thermo Scientific) running in DDA Top10 mode. Based on their hydrophobicity the tryptic peptides were separated using a chromatographic gradient of 60 min with a binary system of buffer A (0.1% formic acid) and buffer B (80% ACN, 0.1% formic acid) with a total flow of 250 nl/min. Separation was achieved on in-house made analytical columns (length: 50 cm, inner diameter: 75 μm) containing 2.7 μm C18 Poroshell EC120 beads (Agilent) heated to 50 °C in a column oven (Sonation). Over a time period of 41 min, Buffer B was linearly increased from 3% to 30% followed by an increase to 50% in 8 min. Finally, buffer B was increased to 95% within 1 min followed by 10 min washing step at 95% B. Full mass spectrometry (MS) spectra (300-1,750 m/z) were recorded with a resolution of 70,000, a maximum injection time of 20 ms and an AGC target of 3e6. In each full MS spectrum, the top 10 most abundant ions were selected for HCD fragmentation (NCE 27) with a quadrupole isolation width of 1.8 m/z and 10 s dynamic exclusion. The MS/MS spectra were then measured with a 35,000 resolution, an injection time of maximum 110 ms and an AGC target of 5e5.

The MS RAW files were then analyzed with MaxQuant suite (version 1.5.3.8) on standard settings. By matching against the human UniProt database the peptides were then identified using the Andromeda scoring algorithm (34). Carbamidomethylation of cysteine was defined as a fixed modification, while methionine oxidation and N-terminal acetylation were variable modifications. The digestion protein was Trypsin/P. A false discovery rate (FDR) < 0.01 was used to identify peptide-spectrum matches and to quantify the proteins. Data processing, statistical analysis, as well as clustering and enrichment analysis were performed in the Perseus software (version 1.6.15.0) (35).

### Immunoblot analysis

Protein samples from co-immunoprecipitation were loaded onto SDS-polyacrylamide gels using SDS-sample buffer, separated by gel-electrophoresis and analysed by immunoblotting. All antibodies were diluted in 50 mM Tris [pH 7.2], 150 mM NaCl with 0.2% Tween-20 and 5% skim milk powder. Antibodies and dilutions are listed in Supplementary Table 1. For visualization, we used Amersham ECL Prime or Select Western Blotting Detection Reagent (GE Healthcare) in combination with the Fusion FX-6 Edge system (Vilber Lourmat).

### Protein structure modelling and visualization

Chimera X Version 1.1 was used to visualize the structure of the ASAP complex (accession number 4A8X on PDB, (20)).

### siRNA-mediated knockdowns

2-3×10^5^ cells were seeded in 6-well plates well and reverse transfected using 2.5 µl Lipofectamine RNAiMAX and a total of 60 pmol of the respective siRNA(s) according to the manufacturer’s instructions. All siRNAs used in this study are listed in Supplementary Table 1.

### RNA extraction, Reverse transcription, endpoint and quantitative RT-PCR

RNA was extracted using different extraction methods. For endpoint or quantitative RT-PCR (RT-qPCR), RNA was extracted using peqGOLD TriFast (VWR Peqlab) or RNA-Solv Reagent (Omega Bio-Tek) following the manufacturer’s instructions for TriFast but using 150 µl 1-bromo-3-chloropropane instead of 200 µl chloroform and eluting the RNA in 20 µl RNase-free water. Reverse Transcription was performed using the GoScript Reverse Transcriptase (Promega), 10 µM VNN-(dT)_20_ primer and 0.5-1 µg of total RNA in a 20 µl reaction volume. RT-PCR and RT-qPCR were performed according to the manufacturer’s protocols using MyTaq™ Red Mix (Bioline/BIOCAT) for RT-PCR and GoTaq qPCR Master Mix (Promega) for RT-qPCR. All primers used in this study are listed in Supplementary Table 1.

### RNA-Sequencing and computational analysis

HTO or HEK 293 cells and the indicated rescue cell lines were treated with siRNA as described above. HTO sets were harvested using TriFast, the HEK 293 set was harvested using RNA Solv Reagent. Extraction of total RNA was performed with DIRECTzol Miniprep Kit (Zymo Research), according to the manufacturer’s instructions.

For each sample, three biological replicates were analyzed. The Spike-In Control Mix (SIRV Set1 SKU: 025.03, Lexogen), which enables performance assessment by providing a set of external RNA controls, was added to the total RNA, as listed in Supplementary Table 2. The Spike-Ins were used for quality control purposes, but not used for the final analysis of differential gene expression (DGE), differential transcript usage (DTU) or alternative splicing (AS). The cDNA library was prepared using the TruSeq Stranded Total RNA kit (Illumina). Library preparation involved the removal of ribosomal RNA using biotinylated target-specific oligos combined with Ribo-Zero Gold rRNA removal beads from 1 µg total RNA input. the Ribo-Zero Human/Mouse/Rat kit depleted cytoplasmic and mitochondrial rRNA from the samples. Following a purification step, the remaining RNA was fragmented and cleaved. The first strand cDNA was synthesized using reverse transcriptase and random primers. Subsequently, the second strand cDNA synthesis was performed using DNA Polymerase I and RNase H. To the resulting double-stranded cDNA, a single ‘A’ base was added and the adapters were ligated. After this, the cDNA was purified and amplified with PCR, followed by library validation and quantification on the TapeStation (Agilent). Equimolar amounts of library were pooled and quantified using the Peqlab KAPA Library Quantification Kit 587 and the Applied Biosystems 7900HT Sequence Detection System. Sequencing was performed on an Illumina NovaSeq6000 sequencing instrument with an PE100 protocol.

The resulting reads were aligned against the human genome (version 38, GENCODE release 33 transcript annotations (36), supplemented with SIRVomeERCCome annotations from Lexogen; (obtained from https://www.lexogen.com/sirvs/download/) using the STAR read aligner (version 2.7.3a) (37). Salmon (version 1.3.0) (38) was used to compute estimates for transcript abundance with a decoy-aware transcriptome. Transcript abundances were imported, followed by differential gene expression analysis using the DESeq2 (39) R package (version 1.28.1) with the significance thresholds |log2FoldChange| > 1 and adjusted p-value (padj) < 0.05. Differential splicing was detected with LeafCutter (version 0.2.9) (40) with the significance thresholds |deltaPSI| > 0.1 and padj < 0.001. Alternatively, rMATS (version 4.1.1, (41)) with novel splice site detection was used to identify alternative splicing (AS) classes, followed by analysis using maser (version 1.8.0) and significance thresholds |deltaPSI| > 0.2 and padj < 0.01.

Differential transcript usage was computed with IsoformSwitchAnalyzeR (ISAR, version 1.10.0) and the DEXSeq method (42-47). Significance thresholds were delta isoform fraction |dIF| > 0.1 and adjusted p-value (isoform_switch_q_value) < 0.05. Intron retention was computed with IRFinder (version 1.2.6, (48)) in FastQ mode and differential intron retention was calculated using DESeq2 with the significance thresholds |log2FoldChange| > 1 and padj < 0.001. Sashimi plots were generated using ggsashimi (version 1.0.0, (49)).

## Results

### RNPS1 plays a minor role in NMD

RNPS1 was shown to regulate multiple types of AS in combination with other ASAP/PSAP components in *D. melanogaster* and human cells (22-25). Furthermore, several studies indicated that RNPS1 is able to activate NMD (29,30,50,51) and more recently RNPS1 was reported to be involved in the recognition of many EJC-dependent NMD substrates (31) (Figure 1A). To investigate the role of RNPS1 in NMD in the context of the EJC, we performed RNA-sequencing (RNA-Seq) analyses of cultured human cells depleted of either RNPS1 or the EJC core factors EIF4A3, MAGOH or RBM8A (Figure 1B). Additionally, we sequenced RNA from stable cell lines expressing siRNA-insensitive RNPS1 or EIF4A3 constructs to rescue the respective siRNA mediated knockdown (KD) (Figure 1B and Supplementary Figure 1A, B). In total, we generated three new RNA-Seq datasets from Flp-In-T-REx-293 (HEK 293) cells and HeLa Tet-Off (HTO) cells and re-analyzed existing RNA-Seq datasets of RNPS1 KD-rescue in HeLa Flp-In-T-REx (HeLa FT; E-MTAB-6564)(25).

**Figure 1:**
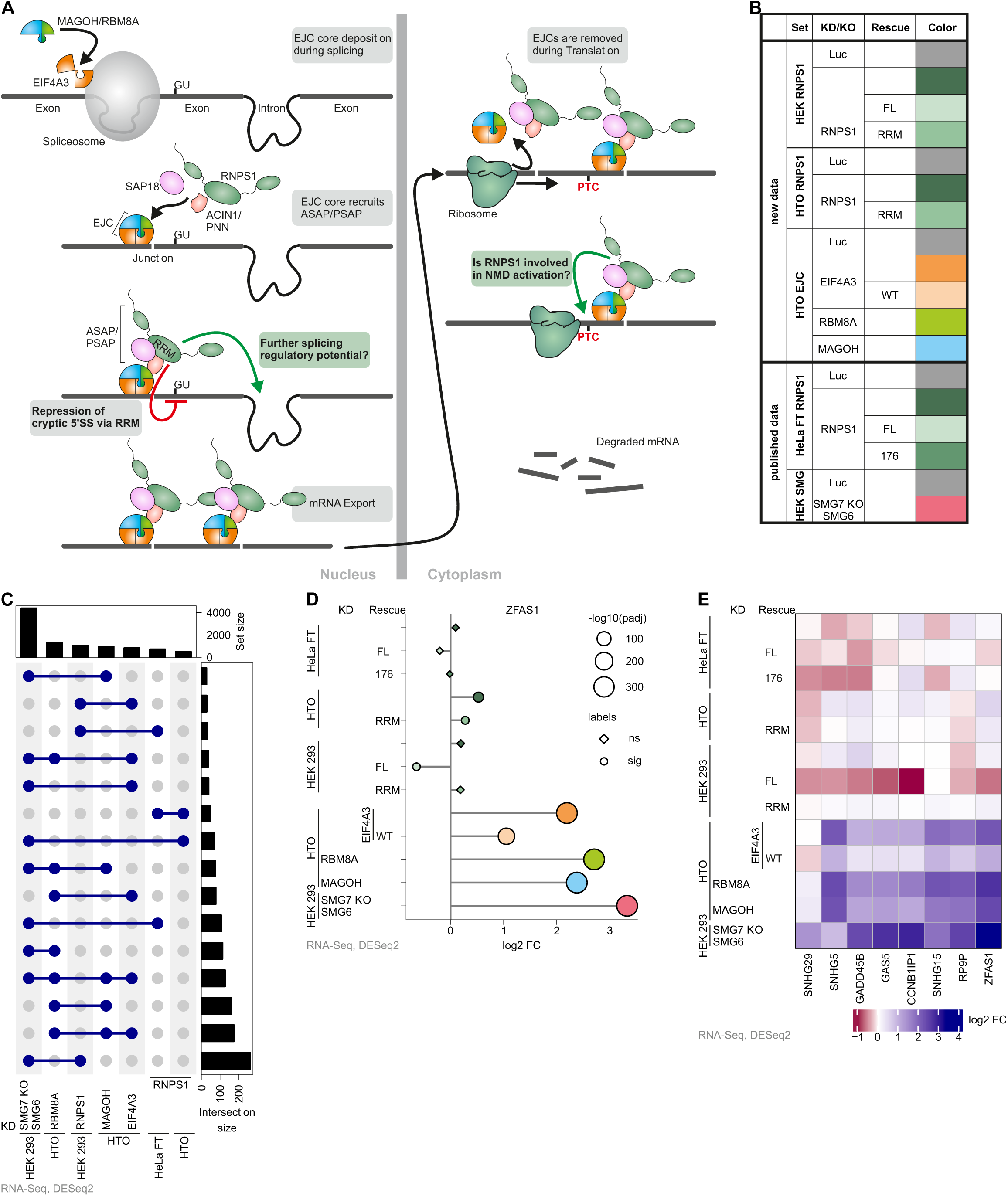
RNPS1 is not required for all EJC-dependent NMD events. **(A)** Schematic depiction of exon junction complex (EJC) deposition on mRNAs during splicing, recruitment of RNPS1-containing ASAP or PSAP complexes to EJCs, alternative splicing regulation (including cryptic 5’ splice site suppression by RNPS1 RRM domain) and NMD activation by RNPS1. Grey boxes indicate established functions and green boxes indicate uncertain functions of RNPS1, which are investigated in this manuscript. **(B)** Overview of published and newly generated RNA-Sequencing (RNA-Seq) data sets, indicating which human cell lines, siRNA-mediated knockdown (KD), CRISPR knockout (KO) and, if applicable, rescue construct was employed. Each condition is assigned to a specific color that is used throughout this manuscript. **(C)** Differential gene expression (DGE) was analyzed using DESeq2 and upregulated genes identified, cutoffs were log2 fold change (log2 FC) > 1 and adjusted p-value (padj) < 0.05. Top 15 intersections between the selected RNA-Seq conditions are depicted in an UpSet plot. P-values were calculated by DESeq2 using a two-sided Wald test and corrected for multiple testing using the Benjamini-Hochberg method. **(D)** DGE analysis of ZFAS1 log2 FCs in the indicated RNA-Seq conditions as compared to the corresponding control. Size depicts the -log10(padj), shape depicts whether the expression change is significant or non-significant (cutoff adjusted p-value (padj) < 0.001). **(E)** Heatmap of log2 fold changes of selected, verified SMG6-SMG7 and EJC-dependent NMD target genes.

Global differential gene expression (DGE) analysis using DESeq2 identified more than 1000 up- or downregulated genes in each RNPS1 KD condition (Supplementary Figure 1C, Supplementary Table 3). The expression of FLAG-emGFP-tagged full-length RNPS1 (RNPS1 FL) or the RRM domain rescued the levels of many genes in HEK 293 and HTO cells. FLAG-tagged RNPS1 FL conferred an almost complete rescue of all events in HeLa FT cells. The rescue with an RNPS1 mutant unable to interact with the ASAP/PSAP complex and the EJC (RNPS1 176) (25), resulted in more mis-regulated genes than the KD alone (Supplementary Figure 1C), which indicates that this mutant exerts a dominant-negative effect.

We hypothesized that if RNPS1 is indeed required for NMD, many NMD-targeted genes should be upregulated upon RNPS1 KD. As a reference for NMD-targeted genes, we used the DGE analysis of a recent RNA-Seq dataset from SMG7 knockout (KO) HEK 293 cells with additional SMG6 KD ((52); E-MTAB-9330), which displayed nearly complete NMD inhibition. The postulated role of RNPS1 as an NMD activator seems to be supported by a substantial overlap of upregulated genes (27-40%) between the three RNPS1 KD conditions and SMG7 KO with SMG6 KD (Figure 1C and Supplementary Figure 1D). However, the overlap between RNPS1 KD with e.g. the KD of the EJC core factor RBM8A was substantially lower (14-28%). When visualizing the extent of differential gene expression of known NMD targets, e.g. snoRNA host genes ZFAS1 and GAS5 (53), we observed only very small effects of RNPS1 KD compared to the strong upregulation upon EJC or SMG6-SMG7 depletion (Figure 1D and Supplementary Figure 1E). Similar trends were observed for other NMD targets (Figure 1E). Of note, the overexpression of RNPS1 FL robustly led to further downregulation of selected known NMD targets, suggesting that elevated RNPS1 levels can enhance NMD (30).

To further characterize the role of RNPS1 in NMD with an orthogonal approach, we analyzed differential transcript usage (DTU) using the IsoformSwitchAnalyzeR package (ISAR) (44). This approach detects upregulated transcripts with annotated PTCs, which indicates NMD inhibition. Depletion of RNPS1 in all three cell lines (HEK 293, HTO and HeLa FT) caused a noticeable upregulation of transcripts bearing a PTC (Figure 2A, Supplementary Table 4), which was quantitatively less pronounced than in SMG7 KO SMG6 KD. Although this in principle supports a role of RNPS1 in the NMD process, we only found a minimal overlap of PTC-containing isoforms between RNPS1 KD and either EJC or SMG6-SMG7 depletion (Figure 2B). In contrast, the overlaps between EJC core factor KDs and SMG6-SMG7 depletion were robust. When plotted against each other, the event strength of differentially used transcripts found in both RBM8A KD and SMG7 KO SMG6 KD showed good correlation (Supplementary Figure 2A). In contrast, there were considerably fewer shared transcripts between RNPS1 KD and SMG7 KO SMG6 KD conditions, which also showed weaker correlation (Supplementary Figure 2B, C). To gain deeper insight into RNPS1’s NMD function, we visualized the RNA-Seq data for bona fide NMD targets, such as SRSF2, where the inclusion of an alternative exon and splicing of an intron in the 3’ untranslated region activates NMD. (54). Both NMD-activating AS events were clearly visible in the EJC- and SMG6-SMG7 depleted conditions, but comparably weak in the RNPS1 KDs (Supplementary Figure 2D). A likely explanation for the minimal overlap of RNPS1-dependent upregulated PTC-containing transcripts are many mRNA isoforms that seemed to be mis-classified as NMD targets by the ISAR algorithm due to unannotated AS events. One example is FAM234B, showing the highest delta isoform fraction (dIF) value in HEK293 RNPS1 KD, in which splicing of an intron in the 3’ untranslated region can produce an NMD isoform (Figure 2C). Accumulation of this PTC-containing isoform is detected in SMG6-SMG7 conditions, whereas in RNPS1- and EJC-depleted cells the skipping of two exons produces an isoform that does not undergo NMD (Figure 2C). Another example is SLC6A6, for which a strictly RNPS1-dependent intron retention event is erroneously counted as upregulation of a PTC-containing isoform (Figure 2D). We found many more cases where our closer inspection revealed that RNPS1 depletion does not lead to the accumulation of NMD-targeted isoforms, but rather to the processing of unannotated and NMD-irrelevant transcripts.

**Figure 2:**
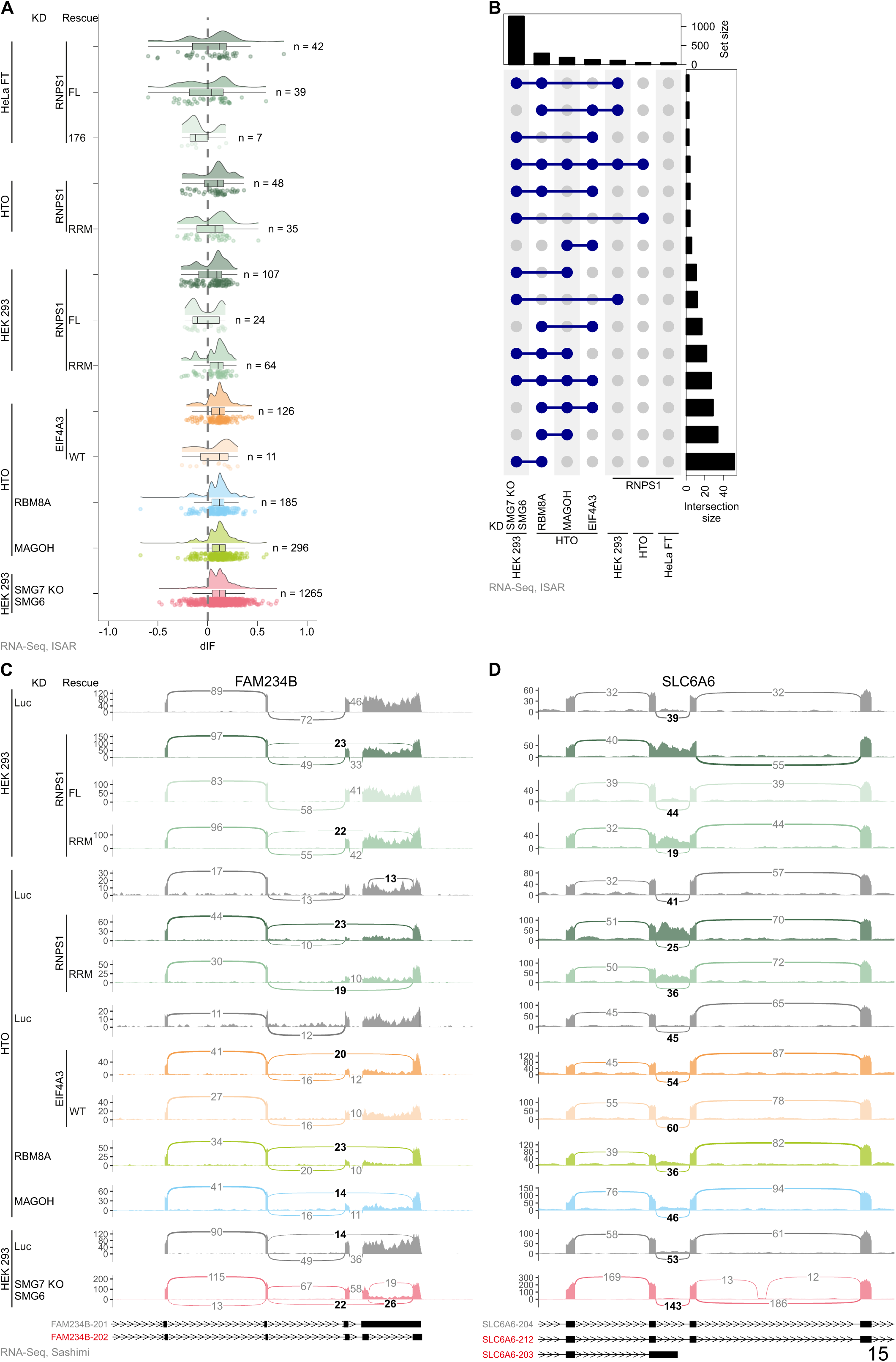
Differential transcript usage analysis reveals RNPS1 main role in alternative splicing rather than NMD activation. **(A)** Raincloud plot depicting the change in isoform fraction (dIF) for GENCODE (release 33) annotated premature translation termination codon (PTC)-containing isoforms (determined via IsoformSwitchAnalyzeR, ISAR) in the indicated RNA-Seq data. Number of individual events with padj < 0.001 cutoff is indicated on the right. P-values were calculated by IsoformSwitchAnalyzeR using a DEXSeq-based test and corrected for multiple testing using the Benjamini-Hochberg method. **(B)** UpSet plot showing the top 15 overlap between the different RNA-Seq conditions with respect to differential transcript usage (DTU). **(C, D)** Sashimi plots show the mean junction coverage of the indicated RNA-Seq data with the canonical and NMD-sensitive isoforms for **(C)** FAM234B and **(D)** SLC6A6 depicted below. NMD-relevant and alternatively spliced junctions are highlighted and NMD isoforms are labeled in red.

In conclusion, at first glance the results from both DGE and DTU analyses seemed to suggest that RNPS1 indeed influences NMD, but in a specific rather than a global way. However, many known NMD targets seemed to be unaffected by RNPS1 depletion, whereas some targets such as ZFAS1 were degraded more efficiently when RNPS1 FL was overexpressed (Figure 1D, E). Especially the DTU in-depth analysis confirmed that the main role of RNPS1 is not in promoting degradation of NMD targets, but in regulating AS (Figure 2C, D).

### Many, but not all RNPS1 dependent alternative splicing events are rescued by RNPS1 RRM expression

Encouraged by the findings of the DTU analysis, we next wanted to further characterize the function of RNPS1 in regulating AS. Previous studies detected hundreds of RNPS1-dependent AS events and experiments with a few individual transcripts demonstrated that the isolated RRM can rescue AS events caused by RNPS1 KD (25). However, the RNPS1 RRM rescue did not normalize splicing of FAM234B or SLC6A6 (Figure 2C, D), suggesting that the RRM cannot perform the same splicing regulatory functions as RNPS1 FL.

To detect transcriptome-wide AS upon loss of RNPS1 or rescue with the RRM domain, we analyzed the RNA-Seq datasets with the intron-centric LeafCutter software (40). Compared to the almost complete restoration of normal splicing by RNPS1 FL, about two-thirds of AS events could not be rescued by the RNPS1 RRM domain (Figure 3A, Supplementary Table 5). However, it proved difficult to define whether an AS event is fully rescued or not since the outcome relied in part on the chosen computational cutoffs. We also observed partially rescued events when we visualized the AS strength as deltaPSI (dPSI) for all AS events found in both the RNPS1 KD and the RRM or FL rescue data (Supplementary Figure 3A, B). These results suggest that the RRM only incompletely rescues RNPS1-dependent alternatively spliced junctions. We validated this partial rescue with selected transcripts (RER1 and FDPS) using RT-PCR and RT-qPCR in both HEK 293 and HTO cells. Both transcripts are alternatively spliced upon RNPS1 KD. RER1 splicing was rescued by RRM expression but in contrast, splicing of FDPS was still impaired in the HEK 293 and HTO cells expressing the RRM (Figure 3B, Supplementary Table 6). This selectivity was also confirmed for two other targets (INTS3 and TAF15), of which INTS3 splicing was rescued, whereas TAF15 was not (Supplementary Figure 3C). Since the RNPS1 RRM is required for ASAP/PSAP assembly, which is also essential for EJC interaction, we speculated that the RRM rescues mainly EJC-dependent splicing events. To this end, we examined exemplary RNPS1-dependent splice events in RNA-Seq data from EJC-protein knockdowns and determined the effects of the RRM rescue. MSTO1 and C5ORF22 are two transcripts with increased AS in the RNPS1 KD, which are also found in EJC KD conditions (Figure 3C, D). While normal splicing of MSTO1 is almost completely restored in RRM-overexpressing cells, the AS event in C5ORF22 remains unchanged in the RRM rescue. Hence, RRM rescue and EJC-dependence do not correlate. These results indicate that, irrespective of the cell line used for the rescue assay and the EJC-dependence of the splice event, the RNPS1 RRM is able to rescue some, but not all AS events in RNPS1 KD.

**Figure 3:**
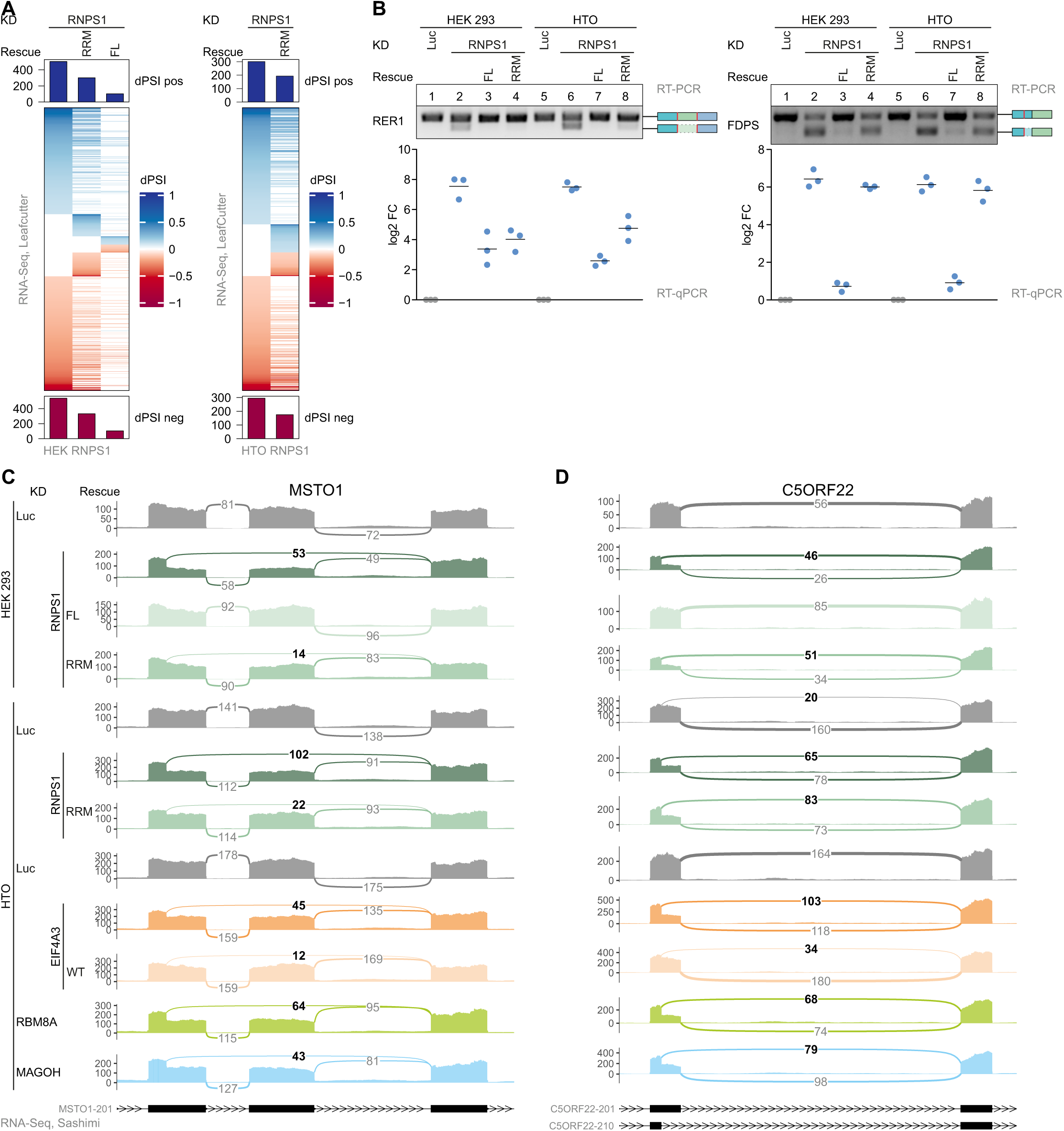
RNPS1 RRM domain partially rescues RNPS1-dependent alternative splicing events. **(A)** DeltaPSI (dPSI) values of ASevents in the indicated RNA-Seq data were calculated with LeafCutter and depicted in a Heatmap (Cutoffs: |dPSI| > 0.1 and padj < 0.001). P-values were calculated by LeafCutter using an asymptotic Chi-squared distribution and corrected for multiple testing using the Benjamini-Hochberg method. **(B)** Comparison of alternatively spliced transcript isoform abundance by RT-PCR and RT-qPCR of RER1 and FDPS in HEK 293 or HTO KD and KD/rescue cells with the resulting PCR product indicated on the right. A representative replicate of the RT-PCRs (n=3) is shown. The log2 FC of RT-qPCRs is calculated as the ratio of alternatively spliced to normally spliced transcript and plotted as datapoints and means. **(C**,**D)** Mean junction coverage in the indicated RNA-Seq condition is depicted as a sashimi plot with the annotated transcript isoforms indicated below and the relevant alternative splice junctions highlighted. Event types are **(C)** Exon skipping (ES) and alternative 5’ splice site (A5SS) usage in MSTO1 and **(D)** A5SS usage in C5ORF22.

### RNPS1 regulates various types of alternative splicing

Previously, RNPS1 was shown to regulate intron retention (IR) in *D. melanogaster* (23,24), but LeafCutter is unable to detect IR events. Hence, we used the IRFinder software to identify RNPS1-regulated retained introns (48). Overall, the RRM rescued more than half of the RNPS1-dependent IR events in HTO and HEK 293 cells (Figure 4A, Supplementary Table 7). However, the splicing of many introns was not rescued at all by the RRM (Figure 4B). The EJC-dependent RFX5 intron 9 retention, for instance, is one of the strongest IR events found in RNPS1 and its splicing was not affected by the RRM (Figure 4C). From a mechanistic point of view, IR is especially interesting, since it represents a seemingly contradictory function of RNPS1: On the one hand, RNPS1 suppresses recursive splicing of cryptic 5’ splice sites, but on the other hand it activates splicing of some introns and thereby represses IR. Therefore, we aimed for a deeper analysis of RFX5 intron 9 splicing. We suspected that splicing of the surrounding introns and subsequent EJC deposition and RNPS1 recruitment reinforces correct RFX5 intron 9 splicing. To test this hypothesis, minigene-reporters in which either one or both introns were deleted were designed and stably transfected into HeLa FT cells (Supplementary Figure 4A Top). RT-PCR of the different reporters shows that RFX5 intron 9 splicing relies on the splicing of both surrounding introns, but mostly on the subsequent intron 10 (Supplementary Figure 4A Bottom). This finding matches our hypothesis that EJCs deposited at the surrounding junctions stimulate splicing of intron 9, similar to what has been described for the PIWI pre-mRNA in *D. melanogaster* (23,24). In a tethering assay, a RFX5 reporter in which intron 10 is replaced by MS2 stem loops was co-transfected with different RNPS1 MS2V5-tagged constructs (Supplementary Figure 4A Top, 4B Top). As seen in the RNA-Seq data, the RNPS1 RRM was unable to rescue RFX5 splicing, even in the tethering assay (Figure 4C, Supplementary Figure 4B Bottom). Interestingly, the RNPS1 176 mutant, which cannot assemble ASAP or PSAP and was unable to rescue most RNPS1-dependent AS events, was able to rescue RFX5 intron 9 splicing when tethered to the mRNA. This indicates that IR repression relies on RNPS1 as the effector molecule and does not require complete ASAP/PSAP complexes or EJC recruitment, once RNPS1 is deposited on the mRNA.

**Figure 4:**
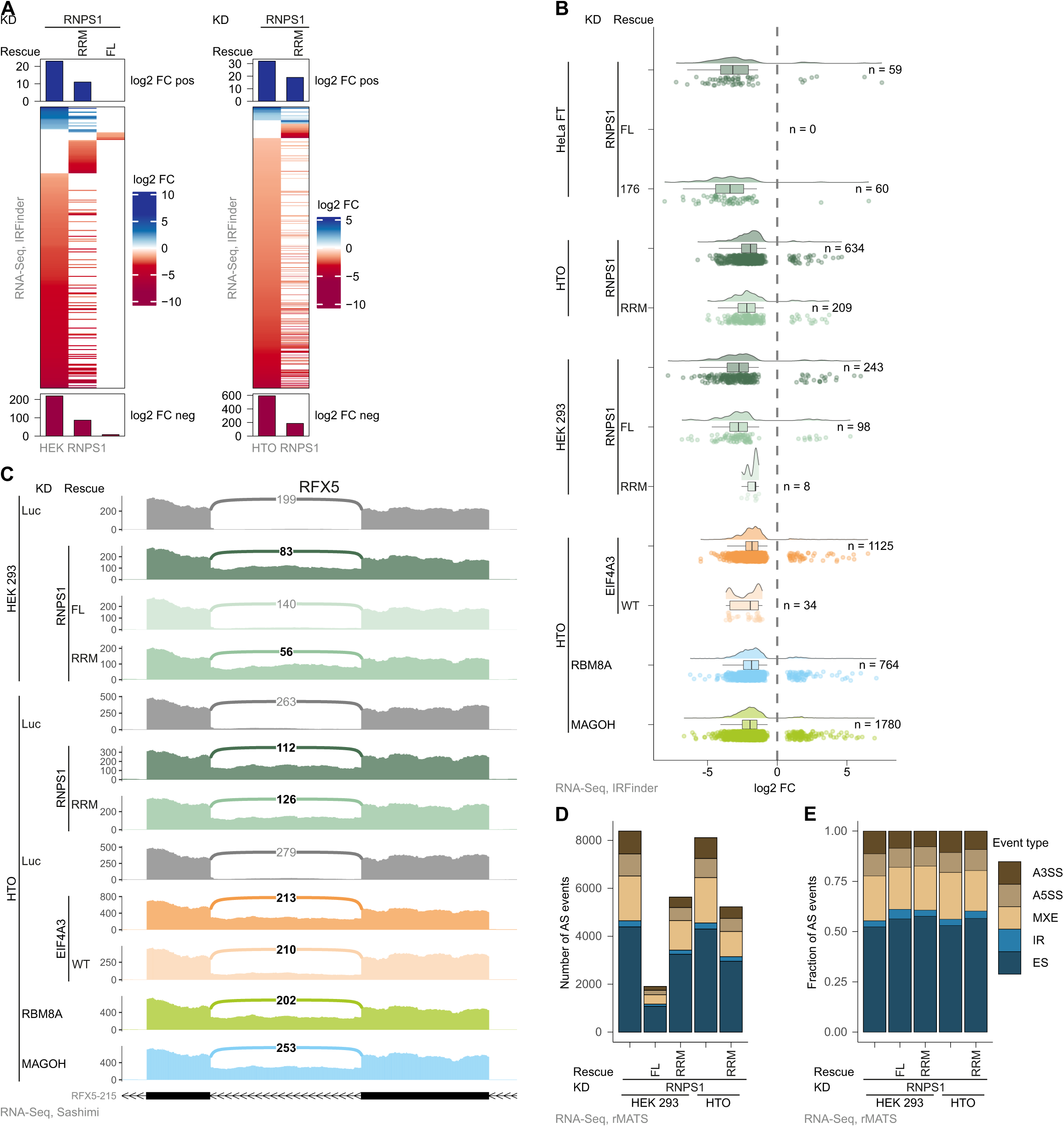
Incomplete rescue with RNPS1 RRM is independent of the alternative splicing type. **(A)** Log2 FCs of retained introns in the different KD and KD/rescue conditions were determined with IRFinder and are depicted in a Heatmap (Cutoffs: |log2 FC| >1 and padj < 0.001). **(B)** The distribution of log2 FC of intron retention (IR) events identified by IRFinder is plotted in a raincloud plot. Number of individual events with padj < 0.001 is indicated on the right. **(C)** RFX5 mean junction coverage in the different RNA-Seq KD and KD/rescue conditions in a sashimi plot with important alternatively spliced junction reads highlighted. **(D, E)** AS event types as detected by rMATS in HEK 293 and HTO RNA-Seq data (Cutoffs: |dPSI| > 0.2 & padj < 0.01). **(D)** Absolute counts, **(E)** Relative fractions.

Although RFX5 correct splicing was not rescued by RNPS1 RRM expression, several IR events were substantially improved, like INTS2 (Supplementary Figure 4C). Therefore, we were wondering whether we can detect discrepancies between RRM rescue of alternative splice sites and of IR. To reveal possible differences, we classified all RNPS1-dependent splice events into categories (exon skipping (ES), alternative 5’ or 3’ splice sites (A5SS/A3SS), exon inclusion (EI) and IR) by using rMATS (41) and determined whether the RRM rescues certain forms of AS. Absolute counts of the various types of AS events and also relative proportions revealed that all types of RNPS1-dependent splicing events were equally well rescued by the RRM (Figure 4D, E, Supplementary Table 8). Taken together, the results of three different bioinformatic analyses show that the RNPS1 KD effects are only partially rescued by the RNPS1 RRM.

### Many splicing associated factors are recruited by the RNPS1 RRM

Despite the incomplete rescue of RNPS1-dependent AS events, our findings suggest that the RRM can regulate different types of AS events, in addition to the previously shown rescue of cryptic 5’ splice sites (25). Therefore, the RNPS1 RRM presumably assembles a splicing-regulatory complex that mediates at least part of the activity of RNPS1 FL. However, neither the mechanism of cryptic 5’ splice site suppression nor the factors interacting with the RRM are known in detail. To characterize the components of this putative splicing-regulatory complex, we set out to identify protein factors that interact with the RNPS1 RRM. We generated HEK 293 cell lines expressing the N-terminally FLAG-emGFP-tagged RRM, confirmed its expression by Western blot (WB) and identified co-purified proteins by mass spectrometry (MS) after FLAG-immunoprecipitation (FLAG-IP; Supplementary Figure 5A). As expected, the RNPS1 RRM efficiently pulled down the three nuclear EJC core components (EIF4A3, RBM8A, MAGOH) and all proteins of the ASAP and PSAP complexes (Figure 5A, B, Supplementary Table 9). Furthermore, RRM-containing complexes contained many factors that are involved in splicing or splicing regulation (Figure 5A). This included spliceosomal or spliceosome-associated proteins (e.g. PRPF6, SNRNP40, U2SURP), many SR proteins (SRSF1, 2, 4, 6, 7 and 9) and other splicing factors, like TRA2A, TRA2B and LUC7L3 (Figure 5A, B). From the various proteins in the RRM interactome, we independently confirmed SRSF7 by Western blotting as a robust interaction partner (Supplementary Figure 5A). We used the Perseus software to cluster proteins enriched in the RRM IP and to identify enriched Gene Ontology (GO) terms for GO biological processes, cellular components, KEGG pathways or PFAM (Supplementary Table 9) (35). Strikingly, the top 10 terms of this analysis are related to mRNA metabolism or splicing (Figure 5C). This is in good agreement with the observation that the RRM domain of RNPS1 is sufficient to regulate specific splicing events. However, it remains to be investigated in more detail, whether all splicing factors interact directly with the RRM or are recruited indirectly via the other proteins of the ASAP or PSAP complex. In conclusion, the RRM of RNPS1 equips the EJC with splicing regulatory abilities by directly or indirectly recruiting splicing-associated factors.

**Figure 5:**
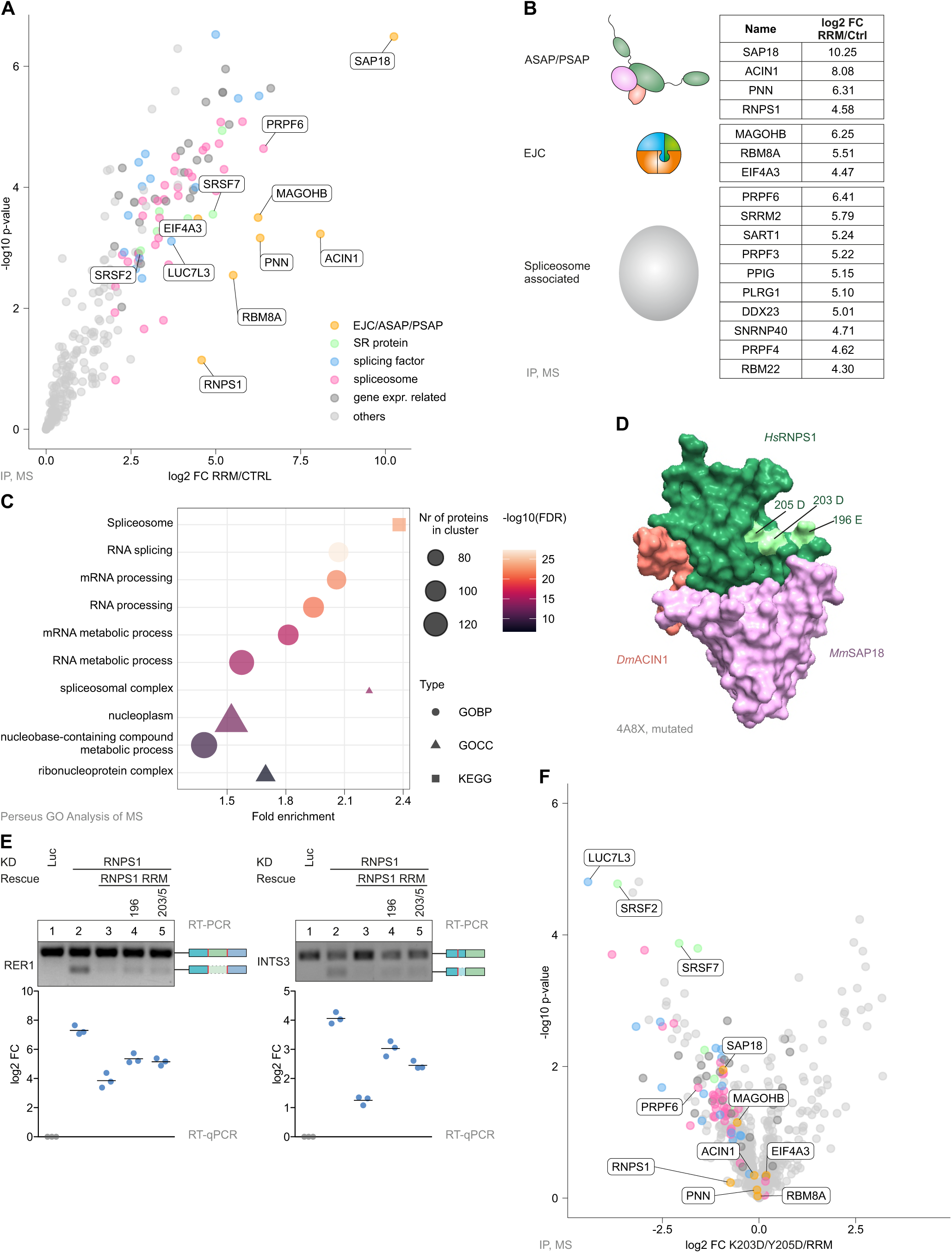
RNPS1 RRM interacts with a broad variety of splicing-regulatory proteins. (**A**) FLAG-RNPS1 RRM construct was overexpressed in HEK 293 cells, followed by FLAG-immunoprecipitation (FLAG-IP) and label-free mass spectrometry (MS). The -log10 p-value of identified proteins is plotted against the log2 FC in a volcano plot (Cutoff: log2 FC ≥0). Proteins identified by MS were manually classified into EJC/ASAP/PSAP proteins, SR-Proteins, splicing factors, gene expression related and others (Supplementary Table 6). (**B**) List of log2 FCs of ASAP/PSAP, EJC and top 10 spliceosome associated proteins found in the RNPS1 RRM MS data. (**C**) Clustered RNPS1 RRM MS results were analyzed for enriched GO terms from GO biological processes (GOBP), GO cellular component (GOCC) and KEGG pathway using Perseus software. -log10 false discovery rate (FDR) of the top 10 terms (according to p-value) are plotted against fold enrichment. (**D**) Structure of the ASAP complex with the indicated RNPS1 mutated residues highlighted in light green (PDB accession number 4A8X, {Murachelli, 2012 #16}). (**E**) RT-PCR and RT-qPCR of RER1 and INTS3 from HEK 293 cells exposed to control or RNPS1 KD and expressing the indicated rescue construct. RT-PCR was performed in triplicates (n=3), one representative replicate is shown and the resulting PCR-product is depicted on the right. For RT-qPCR, the log2 FC of the alternatively spliced transcript to the normal transcript is calculated and plotted as datapoints and means (n=3). (**F**) Label-free MS of FLAG-IP from cells overexpressing the RNPS1 RRM K203D/Y205D construct. The volcano plot shows log2 FC and the corresponding -log10 p-value.

To address the question whether the RNPS1 RRM can directly recruit splicing-regulatory proteins, we used available structural information (20) to mutate potential binding sites on the RRM. To this end, we generated RRM mutants, in which single or multiple surface-exposed amino acids were mutated (Figure 5D). We expected these mutations to allow the formation of ASAP/PSAP complexes, but to disrupt interactions with other splicing effectors. To use these mutants in rescue assays, we integrated them into the genome of HEK 293 cells using the PiggyBac system and induced their expression simultaneously with the siRNA-mediated depletion of endogenous RNPS1. Both mutations, R196E and K203D/Y205D (Figure 5D) reduced the ability of the RRM to rescue the cryptic splicing of 5’ splice sites in RER1 and INTS3, two RNPS1-dependent splice events (Figure 5E, Supplementary Table 6). Interestingly, the two mutants are located close to each other on the surface of the RRM, suggesting that they might interfere with the binding of the same protein(s). These mutants enabled us to identify functionally important interaction partners of the RRM using immunoprecipitation and MS. The analysis of the MS data showed that the mutants were still able to interact equally well with the EJC and ASAP/PSAP complex as the wildtype RRM, which was also validated by WB (Figure 5F, Supplementary Figure 5B, C). This indicates that the ability of the RRM to rescue the AS of RER1 and INTS3 is not reduced because of impaired binding to the ASAP/PSAP complex or the EJC. However, the K203D/Y205D pulled down fewer of the splicing-related factors that are efficiently pulled down by the wildtype RRM (Figure 5F). Although the effect was not quite as pronounced in the R196E pull down, several splicing-regulatory factors exhibited decreased binding to this mutant, too (Supplementary Figure 5C). Two of the most altered factors were SRSF2 and SRSF7, which were far less efficiently pulled down by both mutants.

We therefore conclude that the RRM of RNPS1 is able to regulate splicing by assembling a splicing competent complex, containing inter alia the two SR proteins SRSF2 and SRSF7. The formation of the splice complex is impaired by the mutation of surface patches on the RRM.

### RNPS1 C-terminus mediates interaction with U1 snRNP

The incomplete splicing rescue by the RNPS1 RRM construct clearly demonstrated that other regions of RNPS1 have to play a key role in the regulation of certain splicing events. Therefore, we aimed for an in-depth analysis of the functional domains of RNPS1, for which we followed the well-known domain architecture of RNPS1. As previously mentioned, the RRM domain was shown to be required for ASAP/PSAP assembly (20). The S-Domain, which is located N-terminally of the RRM, was shown to interact with SRP54, while the C-terminal arginine-serine/proline-rich domain (RS/P) interacts with hTra2β (55). No interaction partners are known for the N-terminus. We generated various RNPS1 deletion mutants, lacking either one or two of the RNPS1 domains, stably integrated them into the genome of HEK 293 cells and induced their expression shortly after RNPS1 KD (Figure 6A). Subsequently, we analyzed if the mutants were able to rescue the RNPS1-dependent AS events in FDPS and TAF15. These splicing events were fully rescued by RNPS1 FL and the Del-N variant, but not by any other deletion mutant (Figure 6B). The amount of mis-spliced transcript varied between the RNPS1 variants. The differences in rescue activity indicated that different domains perform partially redundant functions and are not equally important for the activity of RNPS1. However, there were variations in detail and some splice events appeared to be domain-specific, which we could observe, for example, for the AS of FDPS (Figure 6B, C). If rescued with the RNPS1 construct lacking the S-domain, an A5SS in exon 4 was used. When the rescue construct lacked the C-terminus, the same A5SS was combined with an A3SS in the fifth exon of FDPS, resulting in a slightly faster migrating band in the gel. In the RNPS1 KD, approximately 30 % of the FDPS transcripts resulted from only the A5SS while 50 % resulted from alternative 5’ and 3’ splicing and the remaining transcripts were normally spliced. Interestingly, the sashimi plot also shows that the expression of the RRM pushes AS more towards only A5SS usage, similar to the rescue with the Del-S construct (Figure 6B, C).

**Figure 6:**
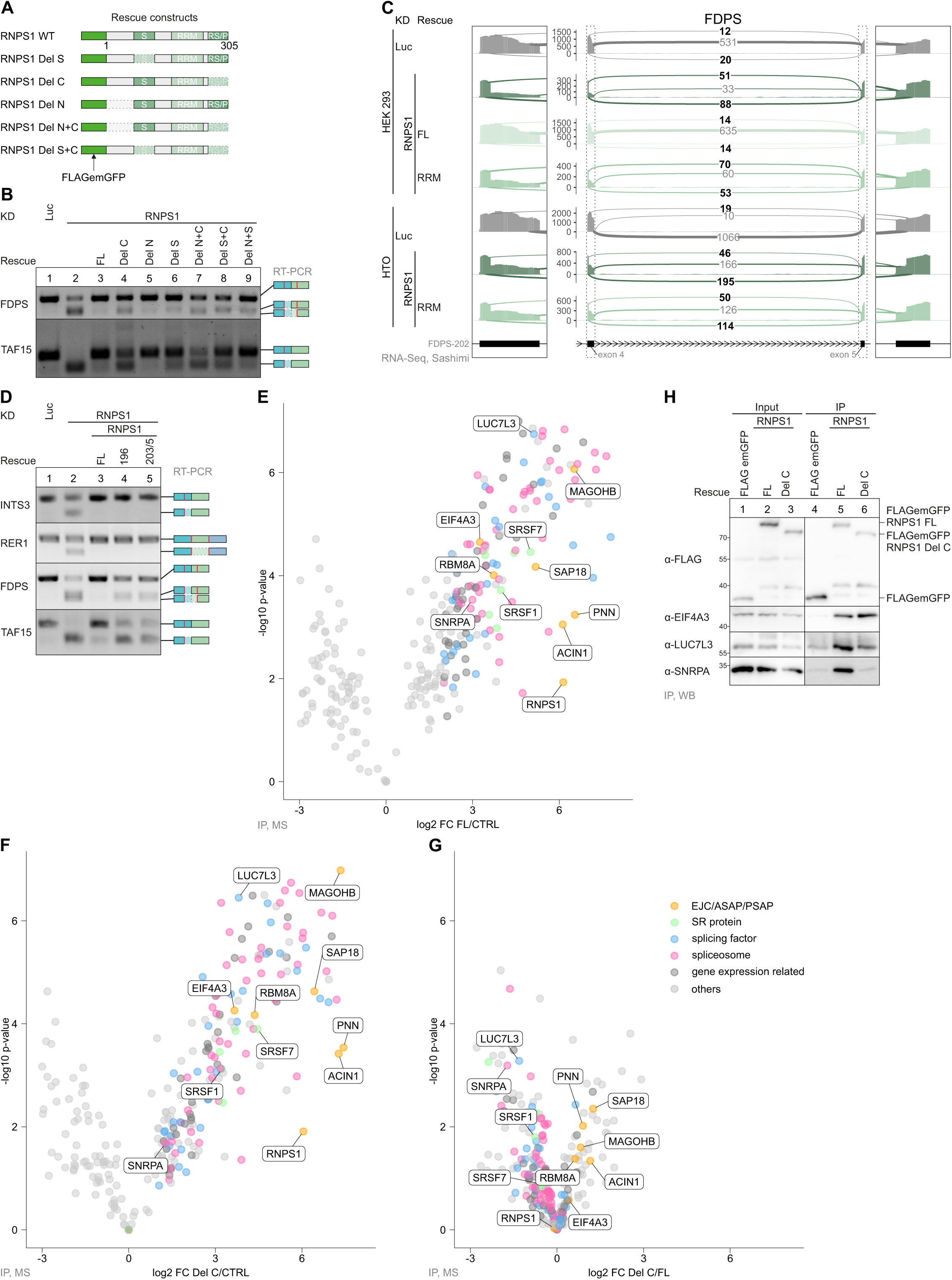
RNPS1 C-terminus is important for regulation of specific alternative splicing events. (**A**) Schematic representation of RNPS1 rescue constructs and domain deletions. (**B**) RT-PCR of FDPS and TAF15 AS in HEK 293 cells after control or RNPS1 KD with the rescue constructs depicted in (A). One representative replicate is shown (n=3). (**C**) FDPS RNA-Seq mean junction coverage is shown as sashimi plot with alternatively spliced junction reads highlighted. (**D**) RT-PCR of INTS3, RER1, FDPS and TAF15 was performed in triplicates (n=3) in the indicated KDs and KD/rescues. One representative replicate is shown with the resulting PCR product depicted on the right. (**E, F, G**) The -log10 p-value of FLAG-IP MS plotted against log2 FC in a volcano plot. For the FLAG-IP, HEK 293 cells overexpressing either a control (CTRL), full-length RNPS1 (FL) or C-terminally shortened RNPS1 (Del-C) were used. Clustering of proteins identified by MS was performed manually into the classes EJC/ASAP/PSAP proteins, SR-Proteins, splicing factors, gene expression related and others (Supplementary Table 7). (**H**) FLAG-IPs of HEK 293 cells overexpressing either a CTRL, RNPS1 FL or RNPS1 Del-C analyzed by Western blot (WB). Antibodies used are shown on the left and a representative replicate is shown (n=3).

These experiments have established that in some transcripts, certain domains of RNPS1 are essential for normal splicing and their deletion cannot be compensated by other domains, for example the RRM alone. Therefore, we wondered to what extent the splicing function mediated by either the RRM or the other regions in RNPS1 substitute each other. Specifically, we asked if the previously generated surface mutations in the RRM (Figure 5) would affect the function of RNPS1 FL and expressed them in RNPS1 depleted HEK 293 cells (Supplementary Figure 6A). The same events in RER1 and INTS3 that were not efficiently rescued by the mutated RRM (Figure 5E) were completely rescued by RNPS1 FL even if its RRM was mutated (Figure 6D, 5E). Unexpectedly, splicing of FDPS and TAF15 could not be rescued by RNPS1 FL carrying the RRM mutations, although FDPS and TAF15 are not rescued by the RRM alone. This leads to the paradoxical observation that in the full-length context, mutations in the RRM seem to affect events that require other domains of RNPS1 for their correct splicing. This suggests that for some splicing events the individual domains of RNPS1 not only perform specific functions, but in some cases also act synergistically together. Even the mutation of RRM surface patches is sufficient to severely disrupt some particularly sensitive events. This suggests that one of the functions of RNPS1 is to locally accumulate a certain concentration of splicing factors by recruiting them via its different domains.

Overall, the deletion of the C-terminus had the strongest effect on the AS events that we analyzed in detail. Therefore, we set out to identify proteins that interact with RNPS1 via its C-terminus. We expressed and immunoprecipitated FLAG-emGFP-tagged RNPS1 FL and the C-terminally (Del-C) shortened version and analyzed the interactome by MS. As expected, we found many splicing factors and components of the spliceosome. However, we did not detect NMD factors (UPF or SMG proteins), supporting our view that RNPS1 has only a minor function in NMD. Notably, several RNPS1-interacting proteins were pulled down less efficiently by RNPS1 Del-C. For example, RNPS1 FL strongly interacted with several U1 snRNP components, whose interaction was significantly reduced with RNPS1 Del-C (Figure 6E, F, G, Supplementary Table 10). The interaction of other splicing factors was also affected by the deletion of the C-terminus. We exemplary confirmed the reduced interaction with the U1 component SNRPA and splicing factor LUC7L3 in a WB (Figure 6H). In contrast, no difference in binding to FL and Del-C RNPS1 was observed for the EJC and the other ASAP-/PSAP components (Figure 6E, F, G, Supplementary Figure 6B). In fact, the pull down of EJC and ASAP/PSAP components even seemed to be slightly improved in the Del-C construct (Figure 6E, G, Supplementary Figure 6B). Overall, our results suggest that the C-terminus of RNPS1 mediates a direct or indirect interaction with the U1 snRNP. This interaction could play a role in the selection of RNPS1-dependent 5’ splice sites.

## Discussion

Although RNPS1 has been the subject of several studies, it remained unclear which of its various functions is most important in the context of the EJC. In this work, we analyze the roles of RNPS1 in NMD and AS regulation and present new molecular details on how AS regulation is mediated by RNPS1. Our analysis of several RNA-Seq datasets suggests that RNPS1 is globally less important for NMD than for example the three core EJC factors EIF4A3, MAGOH or RBM8A. Some NMD targets were upregulated upon RNPS1 depletion, but often only to a low extent (Figure 1D, 1E and Supplementary Figure 1E). Rescue/overexpression of RNPS1, on the other hand, appeared to slightly enhance NMD efficiency. This is in good agreement with previous work in which NMD activation by tethering RNPS1 to reporter mRNAs was shown (50,56). Similarly, the NMD activity of different HeLa cell strains was previously reported to correlate with their RNPS1 expression levels (30). However, some recently reported RNPS1-dependent NMD targets exhibited no consistent response to RNPS1 depletion and were either slightly up- or even slightly downregulated in our RNA-Seq datasets (31). Furthermore, many transcripts that were identified in the DTU analysis as upregulated NMD isoforms in RNPS1 KD turned out - after closer inspection - to be incorrectly classified by ISAR. RNPS1, as previously reported, leads to many new AS events that are not annotated and therefore these new transcript isoforms are apparently mistaken for real NMD isoforms by the ISAR analysis. Altogether, we conclude that RNPS1 is able to increase the NMD efficiency of specific transcripts, which is in good agreement with an NMD-activating function. However, RNPS1 does not interact with NMD factors, raising questions about the mechanism of this effect. Overall, our data leave us with the puzzling observation that the overexpression of RNPS1 stimulates NMD, while the RNPS1 KD has virtually no effect on NMD.

Our finding that many, if not most, of the inspected transcripts identified as RNPS1-dependent NMD targets likely result from AS further emphasizes the importance of splicing regulation by RNPS1. We found previously that the RRM domain is sufficient to regulate some EJC-dependent splicing events (25). Hence, we started our analysis with the initial hypothesis that the RRM domain is sufficient for the regulation of many, if not all RNPS1-dependent splicing events. Unexpectedly, RNA-Seq analyses of RNPS1-depleted cells rescued with the RRM showed that it can only partially replace RNPS1, both quantitatively and qualitatively. Expression of the RRM frequently resulted in incomplete rescue compared to full-length RNPS1. Many other splice events were not rescued at all by the RRM. However, our attempt to classify all RNPS1-dependent splicing events into RRM-rescued and RRM non-rescued did not yield clear results.

During the detailed analyses of the RNA-Seq datasets, we found many of the previously described EJC- and RNPS1-dependent splicing events, for example the use of cryptic 5’ and 3’ splice splices (25-28). In addition, we identified multiple examples of introns, for whose efficient splicing RNPS1 was required. Previous studies in *D. melanogaster* showed that splicing of intron 4 of the PIWI pre-mRNA depends on the deposition of EJCs on exon-exon junctions upstream and downstream (23,24). Our findings demonstrate that this phenomenon is also conserved in human cells. We propose a mechanism similar to that in the fruit fly, namely that the splicing of some weak introns is delayed after the surrounding introns are spliced. Such out-of-order splicing has already been shown for other EJC-dependent splicing events (25). We hypothesize that this splice-activating function of RNPS1 is an important contributor to genome maintenance and prevents inadvertent IR. Mechanistically, multiple EJC-bound RNPS1 proteins appear to cooperate hereby, acting in 3’ and 5’ directions from deposited EJCs.

Our global analysis confirmed that the RRM domain of RNPS1 can rescue different classes or splice events. For this purpose, it interacts with a variety of splicing-related proteins including spliceosomal proteins of different snRNPs. This was an unexpected result, since the main splicing regulatory function of RNPS1 was originally attributed to other domains (55), whereas the RRM seemed to be involved mainly in the formation of the ASAP or PSAP complex (20). However, our results cannot exclude that some of the splicing-related proteins in the RRM interactome are recruited via the other proteins of the ASAP or PSAP complex. PNN, ACIN1 and SAP18 have all been shown to interact with various splicing factors themselves (57-61). Further analyses will therefore be necessary to identify direct interactions between the individual proteins and to disentangle their precise function.

This raises the question if and how other regions of RNPS1 contribute to splicing regulation. To answer this question, we followed previous classifications of the RNPS1 architecture and examined the function of different deletion mutants (55). The deletion of the S-domain affected some splice events, while deletion of the N-terminus had no effect on the examined events. However, we observed the strongest effects with the deletion of the C-terminus and therefore focused our further analyses on the C-terminal region of RNPS1. Our interactome data showed that RNPS1 interacts with U1 snRNP components via its C-terminus (Figure 6G, 7A, Supplementary Figure 7A). Since the U1 snRNP binds to and defines the 5’ splice sites of introns (62), this interaction could be mechanistically involved in the regulation of 5’ splice sites by RNPS1. Interestingly, the suppression of 5’ splice sites was one of the most important functions shown for RNPS1 and the PSAP complex in the context of the EJC (25,26). One seemingly obvious explanation would be that other parts of RNPS1, like the RRM or maybe also the S-domain, repress cryptic 5’ splice sites, while the RNPS1 C-terminus enhances splicing of nearby 5’ splice sites by recruiting the U1 snRNP. Nevertheless, we can only speculate about the exact mechanistic details, and it is also conceivable that the interaction of U1 with the RNPS1 C-terminus prevents cryptic 5’ splicing. Further experiments will be required to unravel the molecular mechanism.

Apart from the interaction with the U1 snRNP, we were able to detect numerous other interactions of RNPS1 and its C-terminus with splice proteins and the spliceosome. These interactions are very interesting from a splicing regulatory point of view and indicate that full-length RNPS1 seems to have a more diverse interactome than the RRM alone, which itself maintains more interactions that its mutants (Figure 7A, B, Supplementary Figure 7B). Considering previous data and the data from this work, we suggest that RNPS1 bridges splicing factors and spliceosomal components to the EJC, thereby recruiting variable splicing competent complexes to the RNA to guide splicing of nearby introns (Figure 7C). RNPS1 seems to act as a “multi-adapter” that binds the U1 snRNP, SR proteins, LUC7L proteins or other splicing factors as required. Due to the amount of different splicing factors that RNPS1 recruits to the pre-mRNA, it can also regulate several different AS events (Figure 7D). The specific way in which RNPS1 acts on each splicing event is determined by the context and the exact position of the EJC. For example, if there are poorly defined introns in its vicinity, RNPS1 stimulates their splicing. In the case of cryptic 5’ splice sites located downstream of RNPS1, it helps to define exonic regions of the mRNA and prevent their re-splicing. The formation of splice-supporting, high-molecular-weight complexes can best explain the role of RNPS1 and could also serve as a model for the mechanism of other multifunctional splicing factors. Moreover, it also fits well with the higher-order mRNP complexes described in the context of nuclear EJC-bound mRNAs (31). Although our model mainly considers the function of RNPS1 in the context of the EJC, it is possible that it can also bind directly to mRNA or is recruited by some of the above proteins to mRNA. This would also explain why not all splicing events regulated by RNPS1 are also EJC-dependent. Interestingly, we had previously observed that the C-terminus can interact nonspecifically with RNA (25). This could reflect its multiple interactions with other RNA-binding proteins, or indicate an intrinsic affinity for RNA.

**Figure 7:**
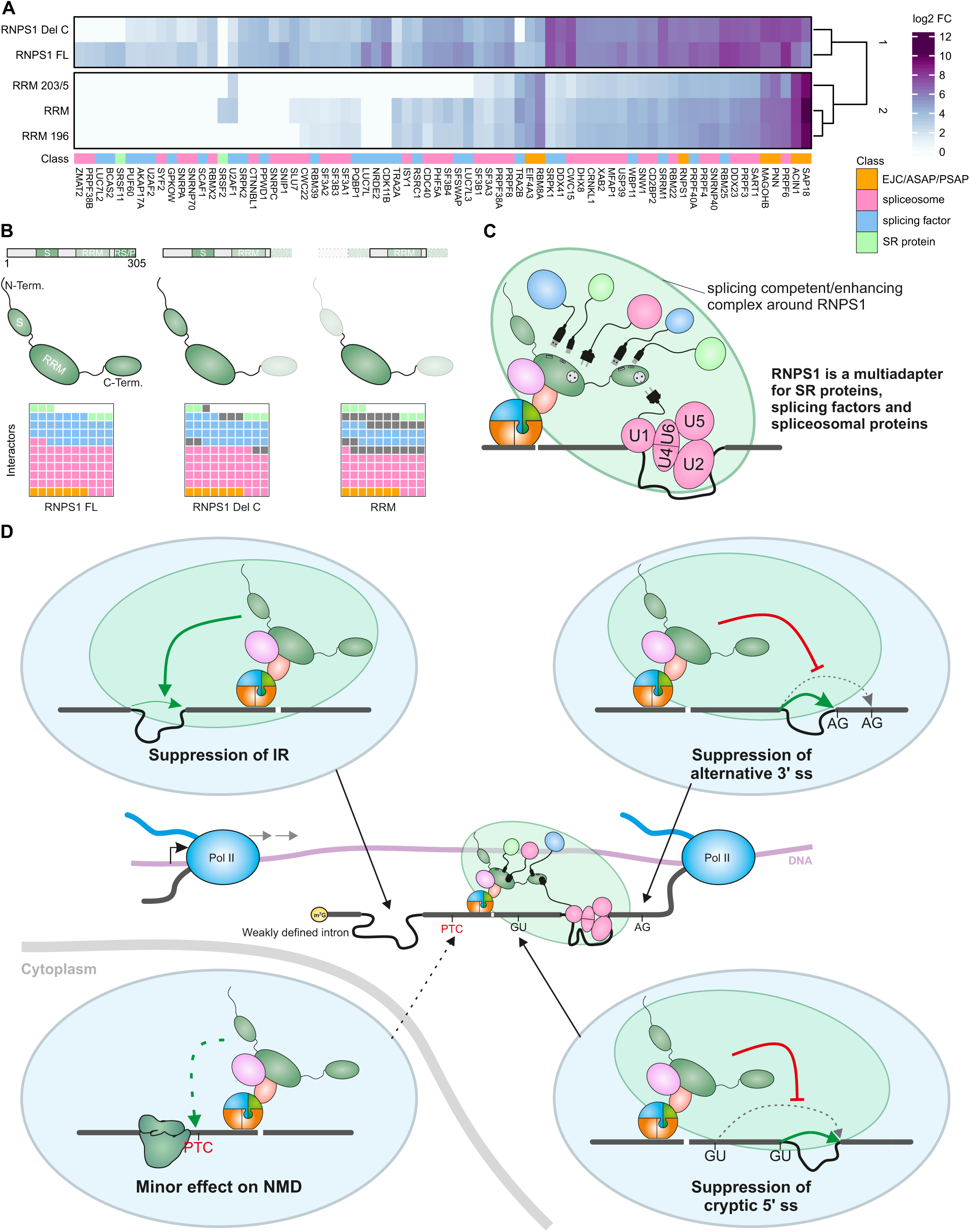
Model for alternative splicing regulation by RNPS1. (**A**) Heatmap showing the log2 FC of RNPS1 interactors compared to the control. Only interactors of the indicated classes are shown. The mean log2 FC for each interacting protein was calculated across all conditions and the sum of the absolute differences of all conditions to this mean had to be > 4. EJC/ASAP/PSAP proteins are shown for comparison. (**B**) Waffle-plots depicting the loss of interaction partners of the indicated classes in the MS of RNPS1 deletion and point mutants. (**C**) RNPS1 interacts with spliceosomal proteins, SR-proteins and other splicing factors with its C-terminus and its RRM. A splicing competent or splicing enhancing complex is formed based on RNPS1. (**D**) Correct splicing of not-well defined introns requires EJC deposition and RNPS1 recruitment via ASAP/PSAP. By assembling a splicing competent or splicing enhancing complex, RNPS1 prevents IR and represses alternative 3’ and alternative 5’ splice sites. NMD is mildly activated if RNPS1 is bound to an mRNA.

RNPS1 is of great interest as a multifunctional splicing protein, because it can either suppress or activate splice sites and enhance the splicing of weak introns, which might seem contradictory at first (Figure 7D). It carries out these functions in conjunction with various other proteins, especially the EJC. This network of interactions allows RNPS1 to regulate a variety of AS events, as it does not rely on its own RNA-binding ability, unlike SR proteins for example. Thus, RNPS1 could be the prototype of flexible, sequence-independent splice regulators, which can be used in regions where no other splicing enhancers can be present due to evolutionary constraints. It will be interesting to find out if other splicing factors can work in a similar way. On the other hand, the interaction of RNPS1 with the EJC needs to be characterized in more detail. So far, we only have some indications how the ASAP complex might interact with the EJC, but more insights will be needed to better understand the 3D structure of EJC-ASAP or EJC-PSAP assemblies. This would also allow us to understand their effect on adjacent splice sites and introns.

## Data availability

RNA-sequencing data generated for this manuscript have been deposited in the ArrayExpress database at EMBL-EBI (www.ebi.ac.uk/arrayexpress) (63) under accession number E-MTAB-10768 [https://www.ebi.ac.uk/arrayexpress/experiments/E-MTAB-10768] for the RNPS1 HTO dataset, accession number E-MTAB-10770 [https://www.ebi.ac.uk/arrayexpress/experiments/E-MTAB-10770] for the RNPS1 HEK 293 dataset and accession number E-MTAB-10770 [https://www.ebi.ac.uk/arrayexpress/experiments/E-MTAB-10770] for the EJC HTO dataset.

Published datasets analysed for this paper include: ArrayExpress accession number E-MTAB-6564 [https://www.ebi.ac.uk/arrayexpress/experiments/E-MTAB-6564] for the RNPS1 HeLa FT dataset (25) and ArrayExpress accession number E-MTAB-9330 [https://www.ebi.ac.uk/arrayexpress/experiments/E-MTAB-9330] for the SMG7 KO and SMG6 KD HEK 293 dataset (52).

The mass spectrometry proteomics data have been deposited to the ProteomeXchange Consortium via the PRIDE (64) partner repository with the dataset identifier PXD027251 [https://www.ebi.ac.uk/pride/archive/projects/PXD027251].

Published protein structure of the ASAP complex was used (PDB: 4A8X [http://doi.org/10.2210/pdb4A8X/pdb]) (20).

All relevant data supporting the key findings of this study are available within the article and its Supplementary Information files or from the corresponding author upon reasonable request.

## Acknowledgements

We thank members of the Gehring lab for discussions and reading of the manuscript. We also thank Marek Franitza and Christian Becker (Cologne Center for Genomics, CCG) for preparing the sequencing libraries and operating the sequencer. NGS analyses were carried out at the production site WGGC Cologne.

## Author contributions

Conceptualization: Lena P. Schlautmann, Volker Boehm, Niels H. Gehring;

Methodology: Lena P. Schlautmann, Volker Boehm, Niels H. Gehring;

Software: Lena P. Schlautmann, Volker Boehm;

Investigation: Lena P. Schlautmann, Volker Boehm, and Jan-Wilm Lackmann;

Resources and Data Curation: Volker Boehm, Janine Altmüller and Jan-Wilm Lackmann;

Writing – Original Draft, Review & Editing: Lena P. Schlautmann, Volker Boehm, Niels H. Gehring;

Visualization: Lena P. Schlautmann and Volker Boehm;

Supervision: Niels H. Gehring;

Funding Acquisition: Christoph Dieterich and Niels H. Gehring;

