## Supplemental Figures for "Exon junction complex-associated multi-adapter RNPS1 nucleates splicing regulatory complexes to maintain transcriptome surveillance"

**This PDF file includes:**

Supplementary Figure 1 to 7

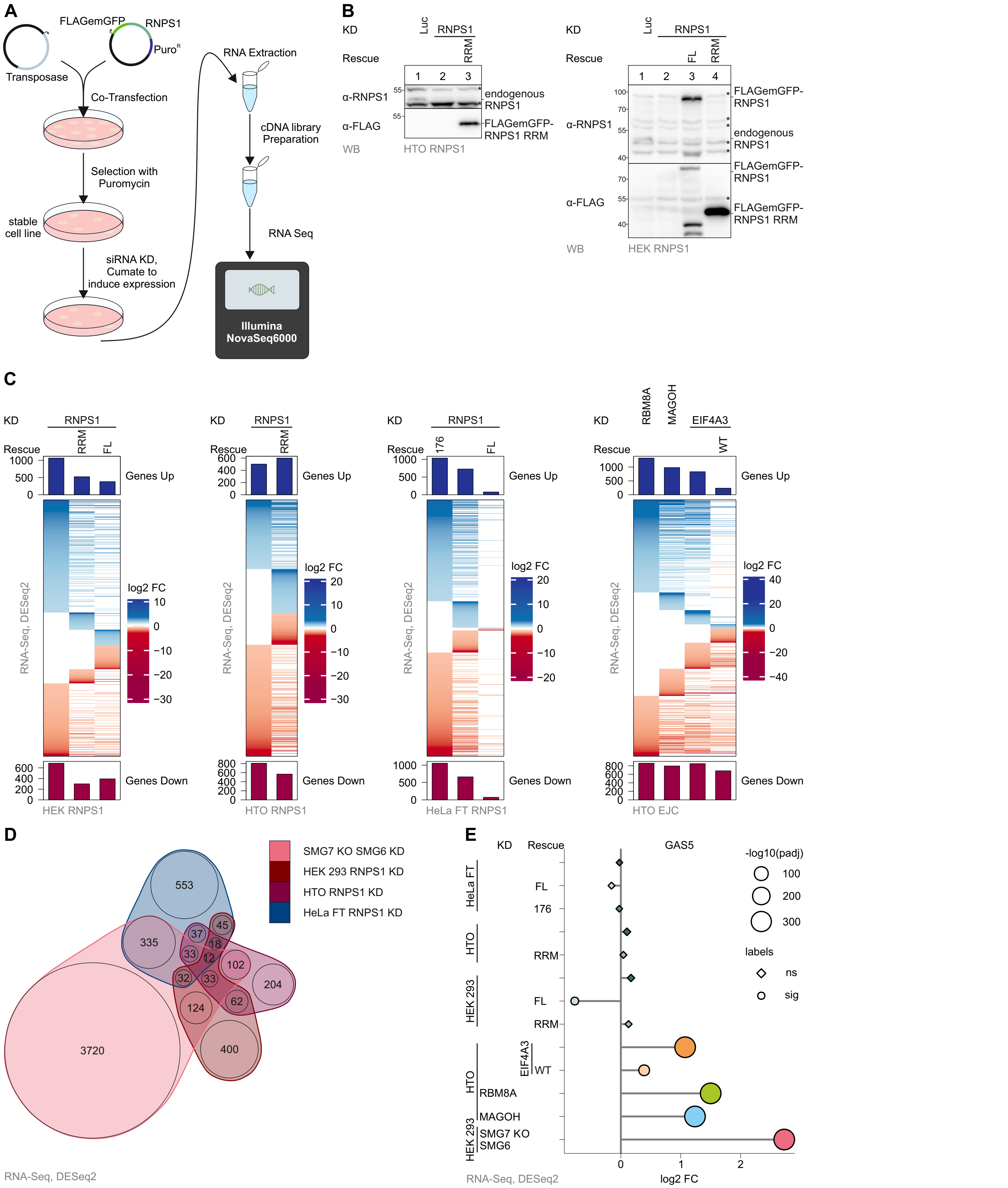

**Supplementary Figure 1: Several genes are mildly upregulated upon RNPS1 depletion.**
**(A)** Workflow of siRNA mediated knockdown (KD) in stable rescue cell lines followed by RNA-Sequencing (RNA-Seq).
**(B)** Western blot (WB) of RNPS1 KD and full-length RNPS1 (FL) or RNPS1 RRM rescue in HTO and HEK 293 cells. Antibodies used are shown on the left and a representative replicate is shown (n=3).
**(C)** Genes up- and downregulated as calculated by DESeq2 for the different RNA-Seq datasets are depicted as a heatmap.
**(D)** nVenn diagram of overlapping upregulated genes in the different RNPS1 KD conditions compared to HEK 293 SMG7 KO SMG6 KD.
**(E)** Differential gene expression (DGE) analysis of GAS5 log2 fold changes (log2 FCs) in the indicated RNA-Seq conditions as compared to the corresponding control. Size depicts the -log10(padj), shape depicts whether the expression change is significant or non-significant (cutoff adjusted p-value (padj) < 0.001).

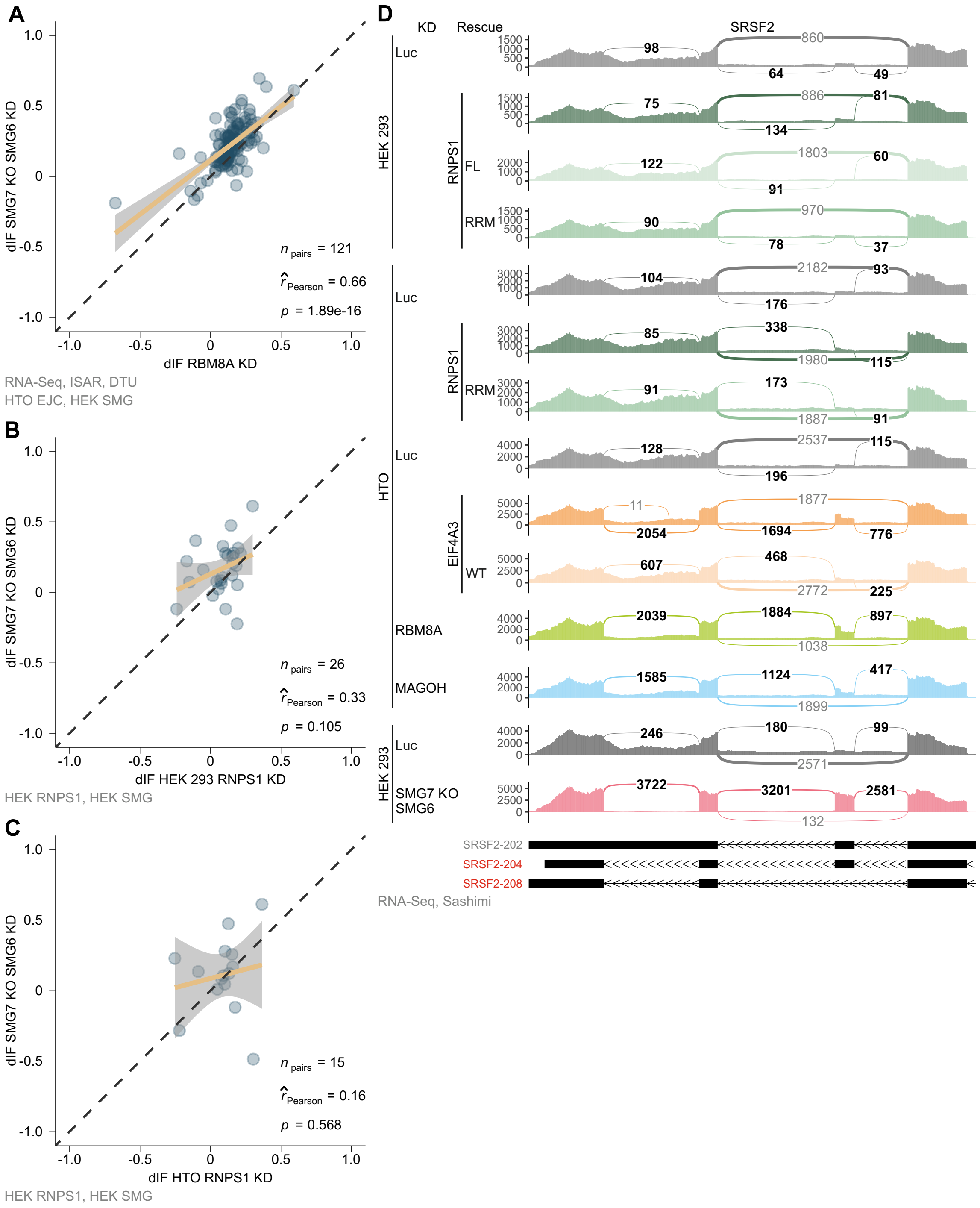


**Supplementary Figure 2: Differential transcript usage does not correlate well between RNPS1 KD and SMG7 and SMG6 depleted cells.**
**(A, B, C)** Scatter plots of differential transcript usage (DTU; cutoff: padj < 0.001) in the indicated RNA-Seq data as calculated by IsoformSwitchAnalyzeR (ISAR). npairs is the number of differentially used transcripts found in both conditions, rPearson is the Pearson correlation coefficient and p is the corresponding p-value. **(A)** RBM8A KD compared to SMG7 KO SMG6 KD, **(B)** RNPS1 KD in HEK 293 cells compared to SMG7 KO SMG6 KD, **(C)** RNPS1 KD in HTO cells compared to SMG7 KO SMG6 KD.
**(D)** Mean junction coverage of the NMD-relevant junctions in SRSF2 in the indicated RNA-Seq KD and KD/rescue conditions is displayed as sashimi plot.

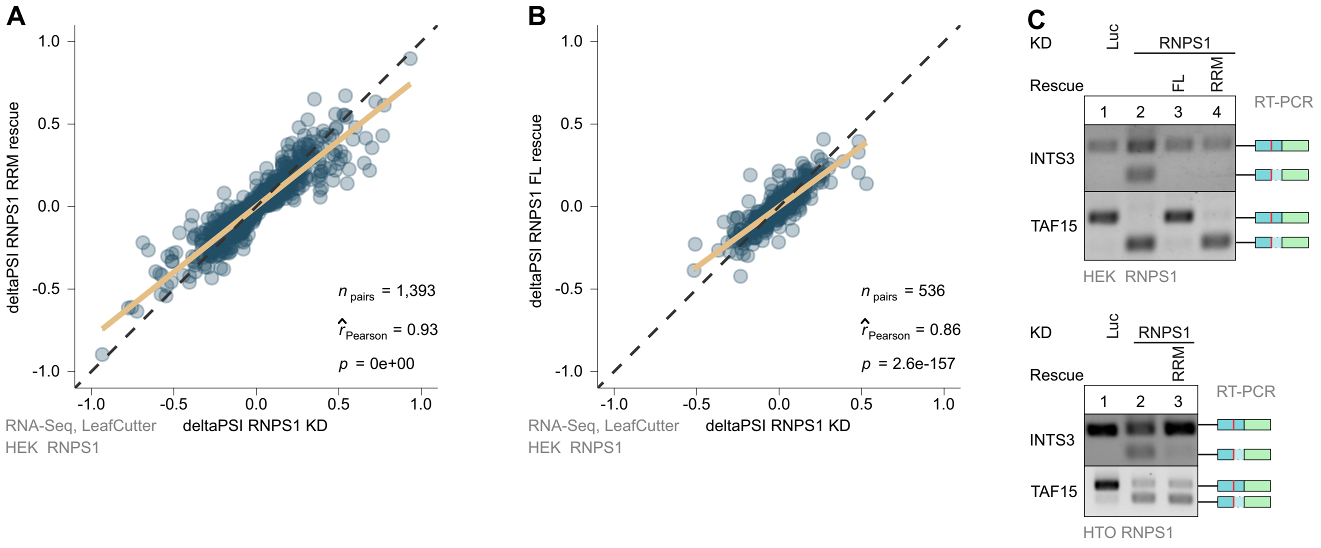


**Supplementary Figure 3: RNPS1 RRM rescue of alternative splicing events is incomplete**

**(A, B)** Scatter plots of alternative splicing (AS) events detected by LeafCutter (Cutoff: padj < 0.001) that are found in both of the indicated conditions in HEK 293 cells. npairs is the number of differentially used transcripts found in both conditions, rPearson is the Pearson correlation coefficient and p is the corresponding p-value. **(A)** RNPS1 RRM rescue compared to RNPS1 KD, **(B)** RNPS1 FL rescue compared to RNPS1 KD.

**(C)** INTS3 and TAF15 AS events in RNPS1 KD and RNPS1 FL or RRM rescue were analyzed by RT-PCR. The resulting AS products are indicated on the right and a representative replicate is shown (n=3).

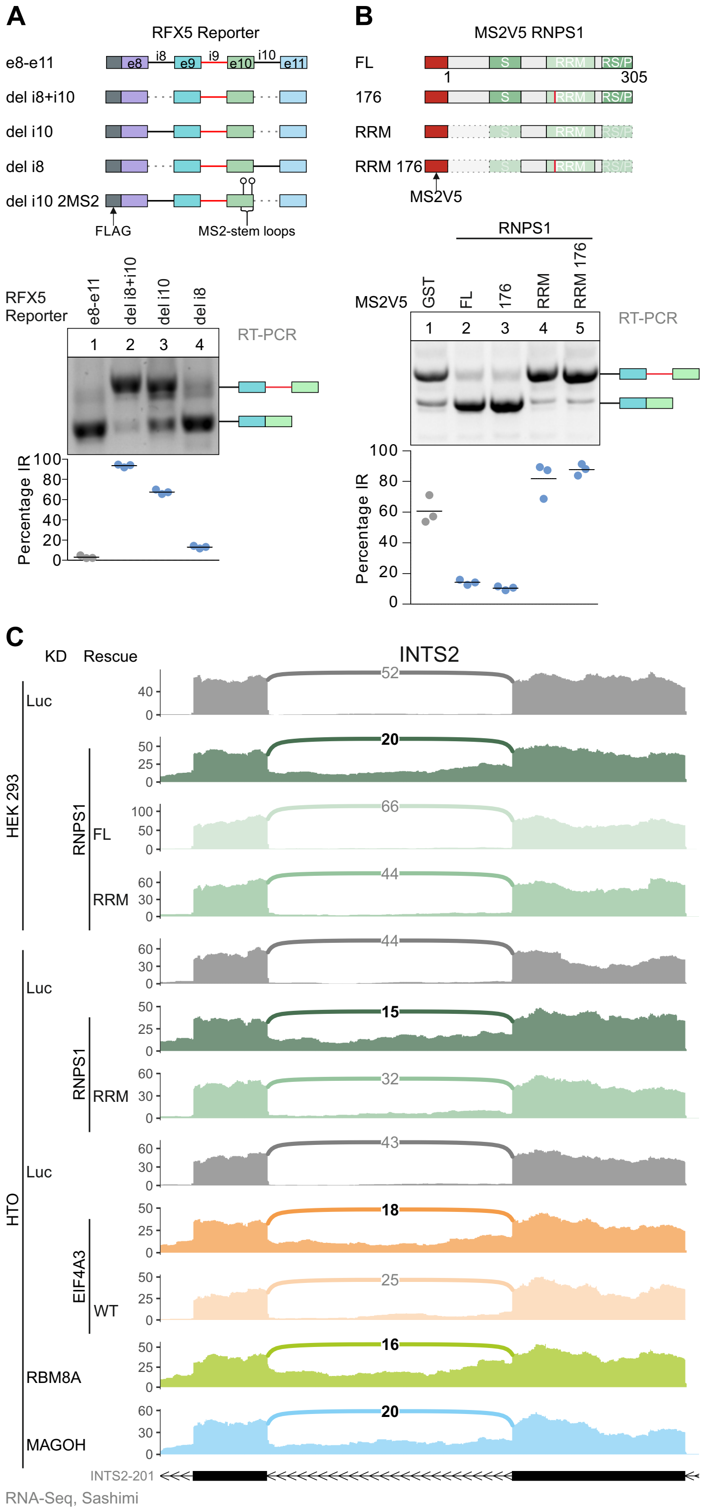


**Supplementary Figure 4: Correct RFX5 intron 9 splicing requires RNPS1 deposition at the subsequent exon-exon junction.**
**(A)** Top: Scheme of RFX5 reporter constructs stably transfected into HeLa FT cells. Bottom: RT-PCR analysis with quantification of intron retention (IR) in the different reporter cell lines, with the resulting PCR-products on the right (n=3).
**(B)** Top: RNPS1 tethering constructs co-transfected with the RFX5 del i10 2MS2 reporter into HTO cells. Bottom: IR was analyzed by RT-PCR followed by quantification, PCR-products are indicated on the right (n=3).
**(C)** Mean junction coverage across the INTS2 potentially retained intron in the different RNA-Seq datasets as Sashimi plot.

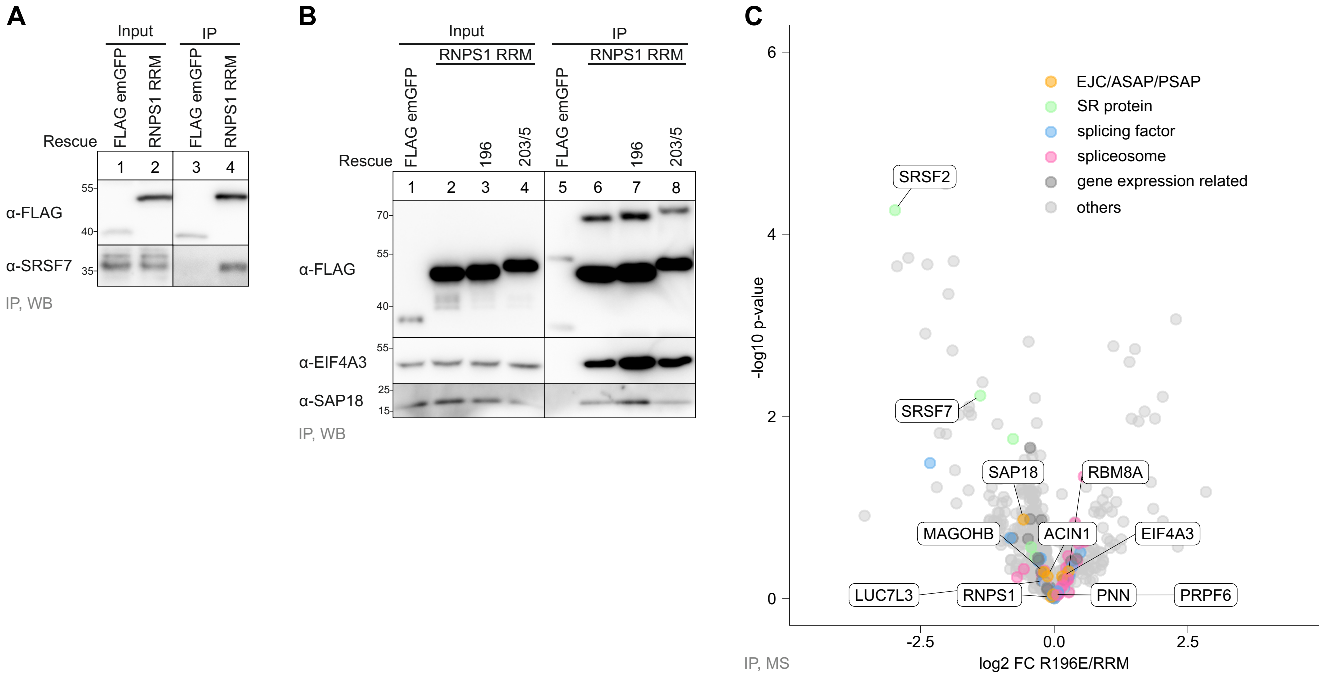


**Supplementary Figure 5: Selected RNPS1 interaction partners confirmed by WB.**
**(A)** WB of FLAG-IP with FLAG emGFP control compared to FLAG-emGFP-tagged RNPS1 RRM (n=3).
**(B)** WB showing the co-immunoprecipitation of the exon junction complex (EJC) component EIF4A3 and ASAP/PSAP component SAP18 by RNPS1 RRM and the RRM 196 and RRM 203/5 mutants (n=3).
**(C)** -log10 p-value of FLAG-IP mass spectrometry (MS) plotted against log2 FC in a volcano plot for RRM 196 mutant compared to the unmutated RRM.

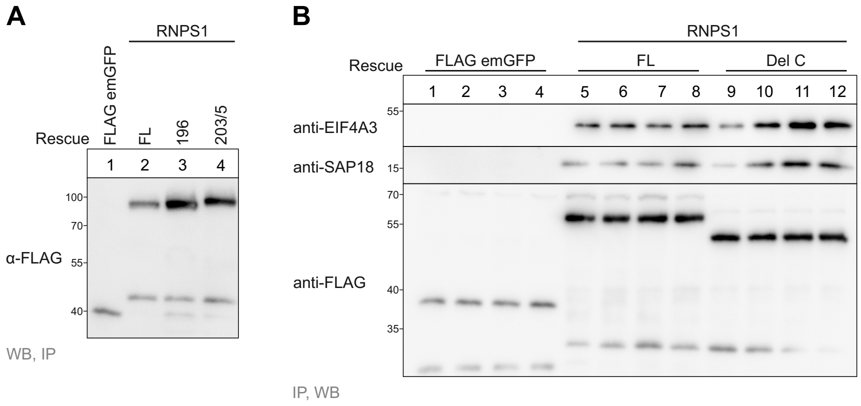


**Supplementary Figure 6: Expression test of RNPS1 mutants and confirmation of EJC and ASAP/PSAP pulldown.**
**(A)** WB with anti-FLAG antibody to detect the expression of RNPS1 point-mutants (n=3).
**(B)** FLAG-IP of control, RNPS1 FL and RNPS1 Del-C is analyzed by WB for co-precipitation of EIF4A3 and SAP18, an EJC or ASAP/PSAP component, respectively (n=3).

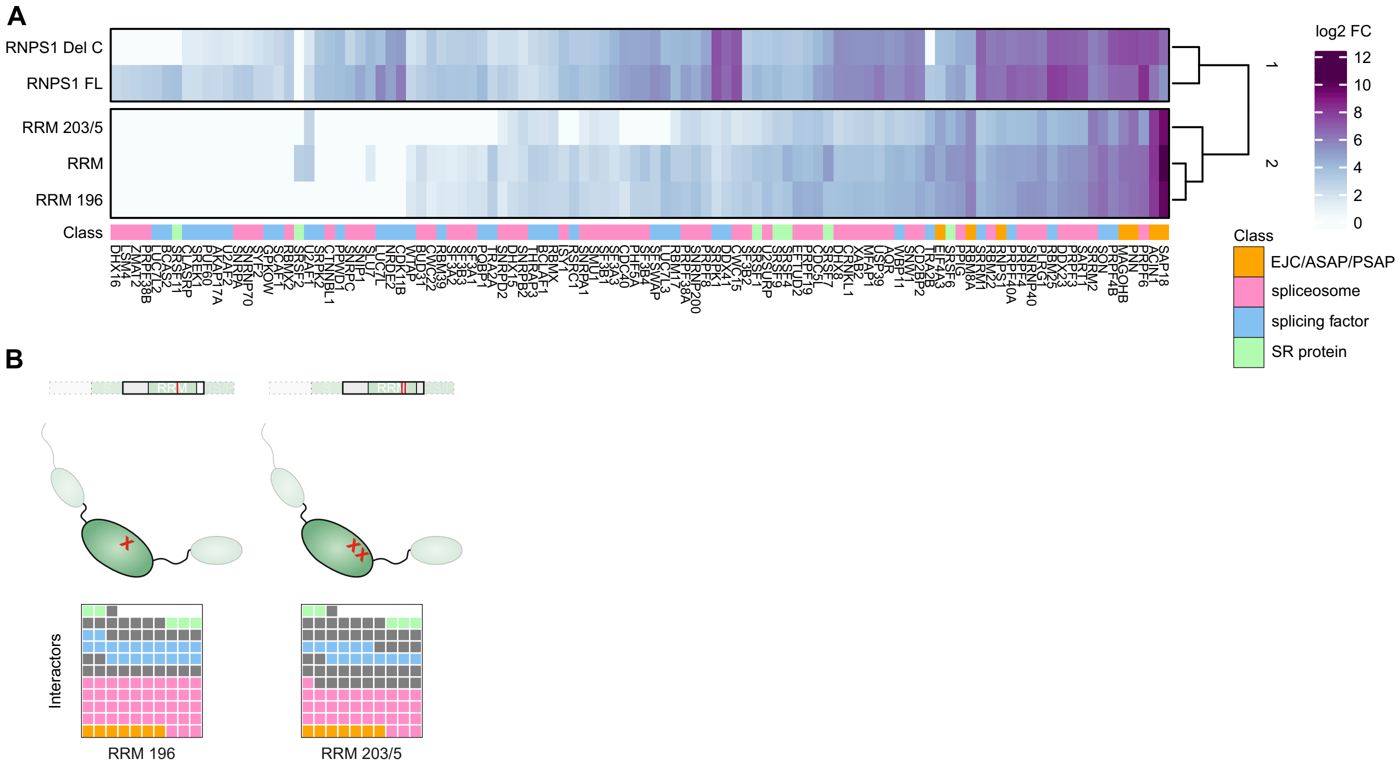

**Supplementary Figure 7: RNPS1 is not required for all EJC-dependent NMD events**
**(A)** Log2 FC of RNPS1 interactors compared to the control are plotted in a heatmap. Only interactors of the indicated classes are shown.
**(B)** The loss of interaction partners pulled down by RNPS1 RRM point mutants depicted as waffle-plots.
